# Mutations beget more mutations – The baseline mutation rate and runaway accumulation

**DOI:** 10.1101/690099

**Authors:** Yongsen Ruan, Bingjie Chen, Qingjian Chen, Haijun Wen, Chung-I Wu

**Affiliations:** State Key Laboratory of Biocontrol, School of Life Sciences, Sun Yat-sen University, Guangzhou 510275, China; CAS Key Laboratory of Genomic and Precision Medicine, Beijing Institute of Genomics, Chinese Academy of Sciences, Beijing 100101, China; Department of Ecology and Evolution, University of Chicago, Chicago, Illinois 60637, USA

**Keywords:** mutation rate, mutation load, mutator, runaway model, cancer risk

## Abstract

There is a sizable literature on mutation rate evolution (Drake 1991; Makova and Li 2002; Lynch 2011; Scally and Durbin 2012; Sung, et al. 2012) but few studies incorporate the recent genomic data from somatic tissues that suggest the operation of mutators. These data show that the mutation burden among cancer samples may vary by several orders of magnitude (Kandoth, et al. 2013; Lawrence, et al. 2013). We now propose a runaway model, applicable to both the germline and the soma, whereby the accumulation of mutator mutations forms a positive-feedback loop. In this loop, any mutator mutation would increase the probability of acquiring the next mutator, thus triggering a runaway escalation in mutation rate. The process can be initiated more readily if there are many weak, rather than a few strong, mutators. Interestingly, even a small increase in the mutation rate at birth could trigger the runaway process, resulting in unfit progeny in slowly reproducing species. In such species, the need to minimize the risk of this uncontrolled accumulation would entail a mutation rate that may seem unnecessarily low. In comparison, species that starts and ends reproduction much sooner do not face the risk and may set the baseline mutation rate higher. The mutation rate would evolve as the generation time changes, therefore explaining many puzzling evolutionary phenomena (Elango, et al. 2006; Thomas, et al. 2010; Langergraber, et al. 2012; Thomas, et al. 2018; Besenbacher, et al. 2019).

## Introduction

Mutational process is fundamental to understanding many evolutionary phenomena. In a lifetime, mutation occurrence is usually treated as a simple Poisson process. This over-simplified assumption seems an inadequate foundation for realistic theories of such phenomena as the germline mutation rate (Kumar 2005; Lynch 2010; Scally and Durbin 2012; Sung, et al. 2012; Wielgoss, et al. 2013; Segurel, et al. 2014; Lynch, et al. 2016), or the evolution of sex (Charlesworth, et al. 2005).

Mutation accumulation in the germline, like that observed in somatic tissues (Alexandrov, et al. 2015; Milholland, et al. 2015; Podolskiy, et al. 2016; Zhang, et al. 2019), is likely a time-inhomogeneous process. In particular, since some mutations are themselves mutators that increase the mutation rate, mutations should beget more mutations thus leading to runaway accumulation. While genes responsible for DNA repair are likely mutators, evidence suggests a much larger number of mutations that may increase the mutation rate in a more subtle manner. For example, any mutation that can influence the chromatin structure or the ion concentration can be a mutator if it can impair the accuracy of DNA polymerase and/or repair enzymes (Schuster-Bockler and Lehner 2012; El Meouche and Dunlop 2018). Unless held back by selection, the first mutator would start a runaway process and eventually greatly elevate the mutation rate. It will be shown that many weak mutators will trigger the runaway more quickly than a few strong mutators, given the same aggregate mutation effect.

In recent years, the accumulation of mutations in somatic tissues (including tumors) has been extensively recorded (Kandoth, et al. 2013; Lawrence, et al. 2013; Martincorena, et al. 2015; Blokzijl, et al. 2016; Ju, et al. 2017; Bae, et al. 2018; Lodato, et al. 2018; Chen, et al. 2019; Yizhak, et al. 2019). The main difference between germline and somatic mutations is that germline mutations are found in all cells of the progeny, whereas somatic mutations affect only local patches of some tissues. The fitness consequences are drastically different. Indeed, somatic mutations experience such weak selection that the process is characterized as “quasi-neutral” (Chen, et al. 2019). Since the runaway process is operative only when it is unchecked by natural selection, somatic mutations are ideal for such an investigation.

In somatic cells, when the runaway mutational process is realized, it often leads to tumorigenesis. Therefore, the end-products of runaway accumulation are different in the germline than in the soma. In the former, they are eliminated by natural selection but, in the latter, they prominently present themselves in tumors. In TCGA (The Cancer Genome Atlas) data, the mutation load varies by more than 1000-fold, even among cases of the same cancer type (Cancer Genome Atlas Research, et al. 2013; Kandoth, et al. 2013; Lawrence, et al. 2013). Hence, the distribution of the mutation load typically exhibits a very long tail among cancer samples (Fig. S1). This highly skewed distribution suggests something akin to a runaway process.

For germline mutations, the mutation rate would be limited by natural selection due to their fitness consequences. Hence, how this limit is set has been a frequent topic in the literature (Drake 1991; Sniegowski, et al. 1997; Lynch 2010; Thomas, et al. 2010; Lynch 2011; Scally and Durbin 2012; Wielgoss, et al. 2013; Segurel, et al. 2014; Jonsson, et al. 2017). Since the mutation rate may not stay constant in a lifetime, it might entail setting the baseline rate low to reduce the likelihood of runaway. There are many other evolutionary implications for avoiding the runaway process, which will be discussed after the theory is presented.

## Results

### I. The runaway mutation accumulation in somatic tissues

While a runaway accumulation of mutations is plausible, it has not been observed presumably because it is too deleterious to the organisms. It is hence expected that natural selection should have stopped the process in the germline. In this regard, somatic mutations would be more revealing as cells that experience runaway accumulation may not die but, instead, transform into tumors.

Fig. 1a shows the mutation load in the coding regions of normal and cancerous colorectal tissues. In the normal tissues, the mutation load is tightly clustered around 40 per exome, whereas the distribution is skewed to the right with a median and mean load of 148 and 503, respectively, in cancerous tissues. The long tail with the load > 5000 is shown in the inset. Note that the mutation numbers in the least loaded 3% of tumor cases fall in the range of the normal tissue. As a result, the highest and lowest numbers among cancer cases differ by 300-fold (~15000/50). Such a long tail is observed in most cancer types (Kandoth, et al. 2013; Lawrence, et al. 2013). We should note that the “tail” is in fact much thinner than the visual impression conveyed in Fig. 1a because cancer patients account for only a small fraction of the population. For example, colorectal cancer patients account for only 2% of the older segment (age > 50) of the general population (https://seer.cancer.gov/). Hence, the load distribution of the entire population would require the red bars be lowered to 2% of the height portrayed in Fig. 1a.

**Fig. 1.**
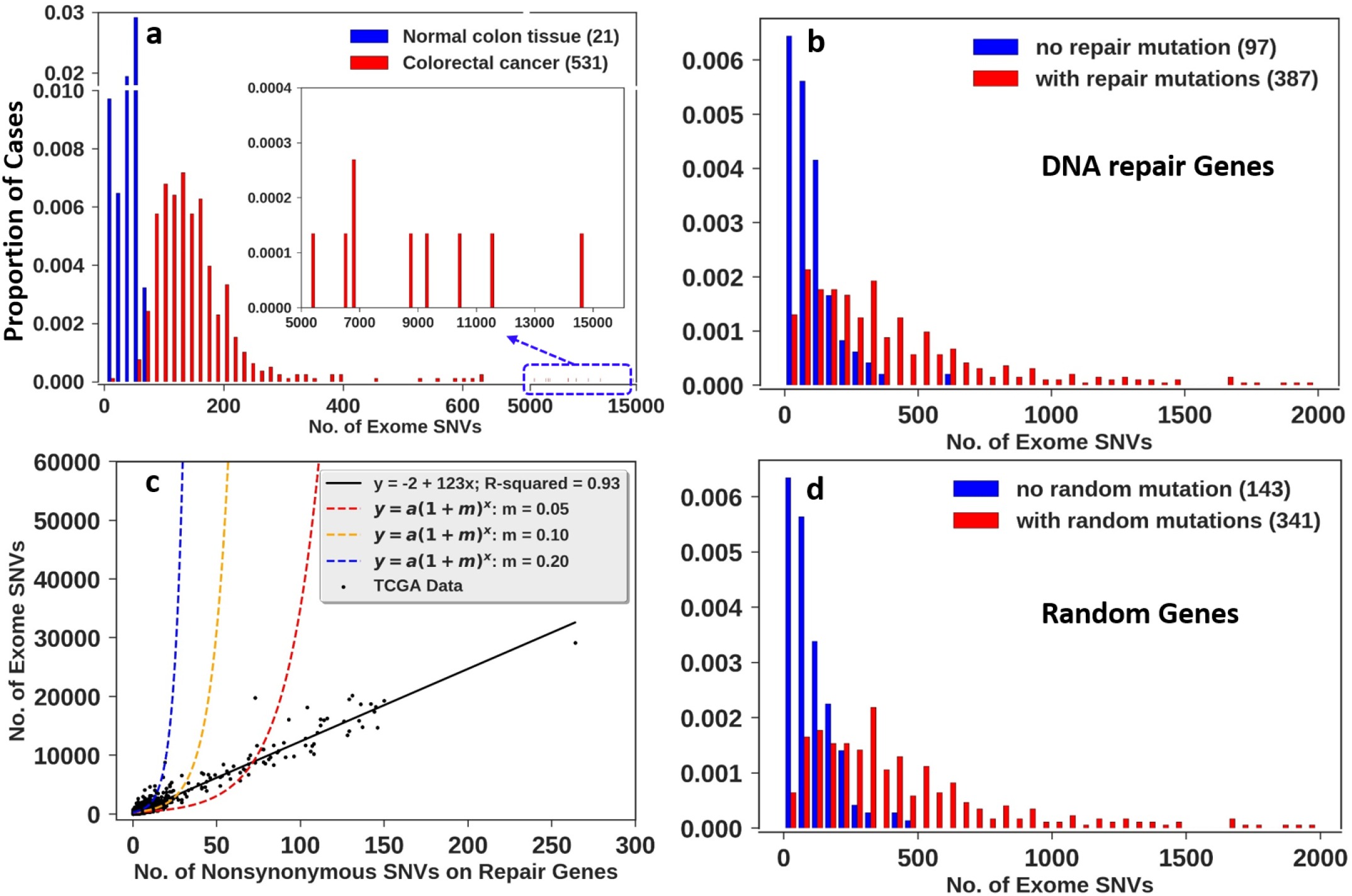
Mutation load distribution. (a) Distributions of mutational load in the normal colon tissue (blue; (Blokzijl, et al. 2016)) and colorectal cancer (red; (Cancer Genome Atlas 2012; Cancer Genome Atlas Research, et al. 2013)). SNVs = single nucleotide variants. The number of cases is shown in the parentheses. The inset displays the tail end of the distribution. (b) Distribution of SNVS in LUAD cancer from TCGA data with and without nonsynonymous mutations in DNA repair genes. Red bars represent cases with at least one nonsynonymous mutation. The number of cases is shown in the parentheses. (c) Mutation load as a function of the number of nonsynonymous mutations in DNA repair genes. Each point represents a patient (9979 patients in total) with the linear regression line shown. The three dashed lines are the models in which each DNA repair mutation increases the mutation rate according to the equation given (see Sec. II). The parameter a = 267 is the mean number of exome SNVs from TCGA data and m is the factor by which a nonsynonymous mutation in DNA repair genes can increase the mutation rate. There is no hint of any increase in mutation rate due to the mutations in DNA repair genes. (d) is the same as (b) but the genes are randomly chosen from the genome with their total length equal to the DNA repair genes. Note that (b) and (d) are nearly identical. They, together with (c), show that the cases with very high mutation load cannot be attributed to the mutations in DNA repair genes.

#### Source of mutators in cancer

If the long tail of the mutation load is caused by mutator mutations, we ask if the mutations in the 177 known DNA repair genes (DRGs) might be the cause (see Methods; (Wood, et al. 2001; Wood, et al. 2005)). In the TCGA data, 4044 of the 9979 cancer samples do not carry mutations in these genes. Fig. 1b separates cases that do or do not have such mutations in red or blue, respectively. Cases with DRG mutations do have a higher mutation load. But the converse is also true: cases with a higher mutation load should have more DRG mutations. We hence randomly choose 207 genes with the same exon length as the 177 DNA repair genes. Indeed, the distribution of nonsynonymous mutations in the randomly-chosen genes is the same as the distribution in the DRGs (Fig. 1b vs. 1d). Hence, the highly loaded cases have exactly the expected number of DRGs mutations. Moreover, if mutations in DRGs really act as mutators, the mutator effect should increase the mutational load exponentially, rather than linearly (Fig. 1c). This will be modeled in Sec. II below. Since the right-skewed long tail is not caused by mutations in the known DRGs, it is plausible that a large number of weak mutators may drive the high mutation rate (Schuster-Bockler and Lehner 2012; El Meouche and Dunlop 2018).

Previous analyses of the mutation load distribution often treat the accumulation as a composite of multiple mutational profiles, referred to as “mutation signatures”. For example, one study identified 33 signatures and suggested five of them to be linearly correlated with the age (Alexandrov, et al. 2015). These clock-like mutations account for ~20% of the total, suggesting that 80% of them accumulate in a time-inhomogeneous manner. Others suggest an exponential increase in the mutation load as a function of age (Milholland, et al. 2015; Podolskiy, et al. 2016; Zhang, et al. 2019). Furthermore, the proportion of clock-like mutations is larger in normal tissues, including early embryos, than in tumors (Blokzijl, et al. 2016; Ju, et al. 2017). These analyses point to the evolution of the mutation rate itself that increases with time. The model in the next section frames such time-dependent mutation rate in mechanistic terms.

### II. The model

In this section, we model mutation accumulation by incorporating a time-dependent mutation rate, *μ*(*t*). A proportion (*ρ*) of mutations is assumed to be mutators that can increase *μ*(*t*) by a fraction *λ*. The mutation accumulation is therefore an inhomogeneous Poisson process. In parallel, we also assume a proportion (r) of mutations to be fitness-altering. In the germline, fitness-altering mutations are usually deleterious but, in the soma, mutations may often be advantageous in the form of cancer driver mutations. Fitness mutations can thus be of both kinds. The remaining fraction [(1-p) (1-r)] of mutations has no phenotypic effect. The process of mutation accumulation lasts for an individual’s lifetime, *T*, during which no recombination happens. *T* follows a truncated normal distribution (μ=70, σ=10, lower=0, upper=120) according to the World Bank Data (https://data.worldbank.org/).

In Fig. 2, examples of mutation accumulation are displayed. All three types of mutations (neutral, fitness-altering and mutator) are shown. Case 1 does not have a mutator mutation; hence, mutations accrue steadily and slowly. In case 2, a large effect mutator leads to a large jump in mutation rate and a quick succession of fitness mutations. In case 3, several small-effect mutators lead to the gradual acceleration of mutation accumulation. In cases 2 and 3, the waiting time between new mutations becomes shorter and shorter, resulting in the accumulation of many fitness-altering mutations.

**Fig. 2.**
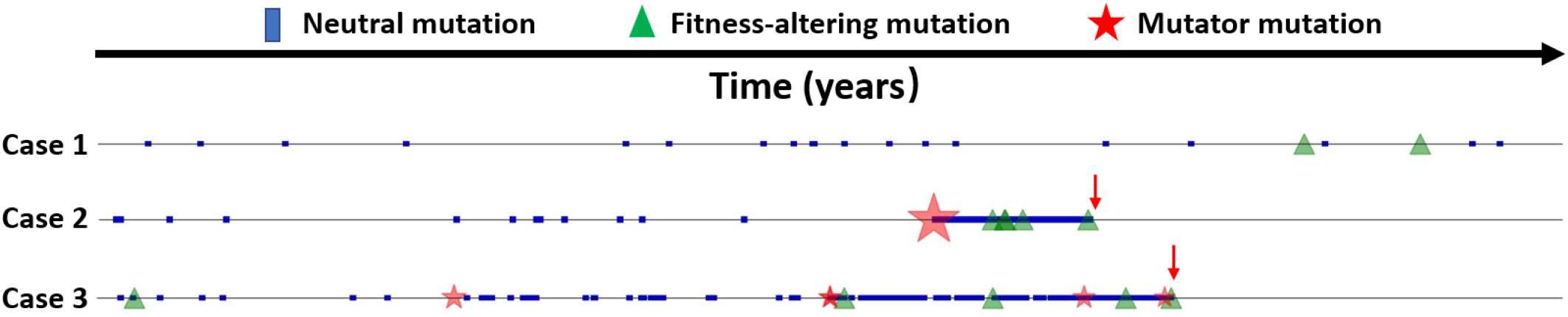
Three examples of mutation accumulation. Three types of mutations are considered: neutral (blue box), fitness-altering (green triangle) and mutators (red star, the size proportional to effect size). Fitness-altering mutations in somatic tissues can lead to cancer when the number reaches a threshold (usually 5). The process lasts for an individual’s lifetime or until reaching the cancerous state.

#### Mutator mutations

Because mutators in the cancer genomic data are not associated with the “strong-effect” DNA repair genes, it seems plausible that mutations affecting chromatin structure, DNA polymerase fidelity, ion concentration in the nucleus, nuclear membrane permeability and numerous other factors may all have a mutator effect, albeit a weak one. In addition, the mutators could be SNVs (single nucleotide variants), CNVs (copy number variations), epigenetic changes or even the expression level of genes (Schuster-Bockler and Lehner 2012; El Meouche and Dunlop 2018). Given that the mutation rate in the human germline at ~10^−9^ /bp /year is very low, a 50% increase in the error rate should be considered weak. (In *E. coli*, strong effect mutators can alter the mutation rate by three orders of magnitude (Loh, et al. 2010)). Nevertheless, if the error rate is multiplicative, 10 weak mutators, each having a 50% effect, would collectively enhance the error rate by 58-fold.

In the model, the baseline mutation rate is *μ*_0_, which is set at 0.33 mutations per exome per year in the soma according to the mutational load of normal tissues (Blokzijl, et al. 2016). The mutation rate changes when a new mutator mutation occurs. (For convenience, we use *μ*^′^ and *μ* and drop the “t”). Thus,

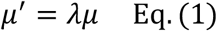

where *λ* is the effect of a new mutator. We assume *ln*(*λ*) = *x* follows the exponential distribution, 1/ *me*^−*x*/*m*^. The mean of the mutator effect is hence 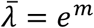 (see Methods). If *m* = 0, the mutator effect *λ* is always equal to 1 and *μ*_*t*_ = *μ*_0_ for the lifetime.

#### Fitness-altering mutations

We now denote the number of fitness-altering mutations by K. The accumulation of these mutations will lead to a phenotypic state, e.g. cancer in the soma or lethality in the progeny (via germline mutations). In the germline, mutations affecting fitness are dominated by deleterious variants (Kimura 1983; Fay, et al. 2001; Eyre-Walker and Keightley 2007; Li 2007) whereas non-neutral mutations are often advantageous in the soma, resulting in the over-proliferation of cells (Fearon and Vogelstein 1990; Hanahan and Weinberg 2011). The soma/germline difference can be seen most clearly in the Ka/Ks ratio, where Ka is the number of nonsynonymous substitutions per site and Ks – synonymous substitutions per site (Li, et al. 1985). For germline mutations, the Ka/Ks ratios range between 0.05 and 0.3 from a wide range of taxa (vertebrates, insects, nematodes, and plants), suggesting 70% to 95% of nonsynonymous mutations are removed by natural selection. In contrast, the Ka/Ks ratio ranges between 0.9 and 1.15 in normal and cancer tissues (Chen, et al. 2019), thus suggesting the prevalence of advantageous mutations in the soma. Due to the relatively weak influence of negative selection in the soma, the runaway process is prominent in cancer samples. We shall present the analysis of somatic mutations first.

1) Fitness-altering mutations in the soma leading to tumorigenesis - A mutation is assumed to affect fitness with probability r. Based on the analysis of 299 cancer driver genes (Bailey, et al. 2018), we set r at 0.04 (See Methods; (Wu, et al. 2016)). The number of genetic events needed to start tumorigenesis has been estimated to be 5 – 10 (Fearon and Vogelstein 1990; Hanahan and Weinberg 2011). Mutations underlying these events are often referred to as “cancer drivers” which, by definition, are advantageous mutations. The cancer risk (R_0_) is the probability of developing cancer in one’s lifetime. Hence, in the deterministic runaway model,

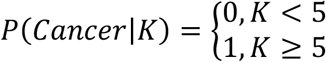

In the probabilistic runaway model, the potential function is:

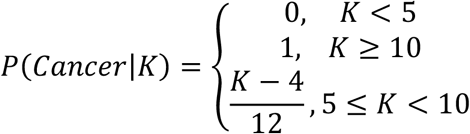

Beyond driver and neutral (passenger) mutations, the existence of deleterious mutations in tumors has been controversial (Wu, et al. 2016; Martincorena, et al. 2017; Zapata, et al. 2018). Such mutations presumably could slow down cancer evolution and should be relevant only when data on non-cancerous clonal expansions are included (see Chen, Ruan et al., unpublished results).

2) Fitness-altering mutations in the germline – We assume that non-neutral mutations occur among 1% (r = 0.01) or 5% (r = 0.05) of the coding mutations on the basis of human polymorphism data (Fay, et al. 2001; Eyre-Walker, et al. 2006). For the purpose of this study, we need to estimate the proportion of mutations under strong negative selection. In the germline, the mutators and the variants that affect fitness are likely to be unlinked or at most loosely linked. Hence, the fitness of the mutators would be affected for only a few generations. For that reason, only fitness reduction approaching lethality is relevant in this context and polymorphism data can distinguish between strongly and weakly deleterious mutations. Fay et al. estimated the proportion, as well as the selective coefficients, of deleterious mutations from the human polymorphism data (Fay, et al. 2001). Their estimates suggest that the strongly selected class (*f*_*2*_, in their notation) is not smaller than 1% - 5% of all coding mutations (see Methods). We assume that these mutations would cause lethality in a combination of two or more. Furthermore, recombination operates in the germline and we assume that the deleterious mutations and the mutators are unlinked.

#### Approximate analytical solutions

While we rely on extensive simulations, approximate analytical solutions have also been obtained as summarized below (see Supplement for details):

The expected number of mutations at age t is given by

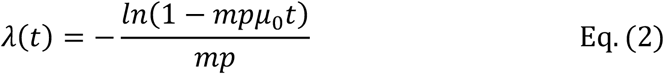

The probability of an individual with *x* mutations at age t is

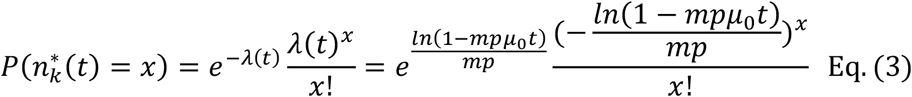

Similarly, we can obtain the probability of having x mutator mutations as

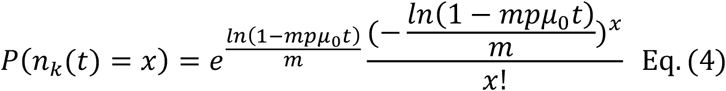

The probability of having x fitness-altering mutations is

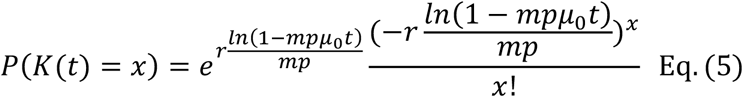

In the deterministic runaway model, the risk at age t is given by

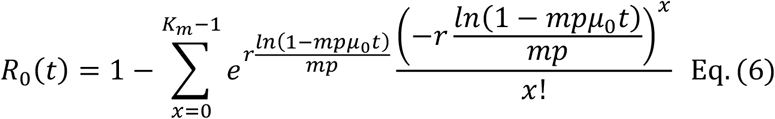

The comparisons between simulations and the analytical solutions are given in Fig. S3 and Fig. S4. The disparity increases with the mutator strength due to the approximation based on the mean mutator effect (see Supplement).

### III. Cause of the runaway process: A few strong or many weak mutators?

To trigger the runaway process, the mutator strength 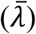 and the initial baseline mutation rate (*μ*_0_) are both crucial. In this section, we analyze the mutator strength (strong but infrequent vs. weak but common mutators) by varying p and 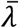 but keeping 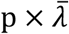 constant (at 0.12). The results are shown in Fig. 3 for the deterministic model. The probabilistic model yields qualitatively similar results as shown in Fig. S2.

**Fig. 3.**
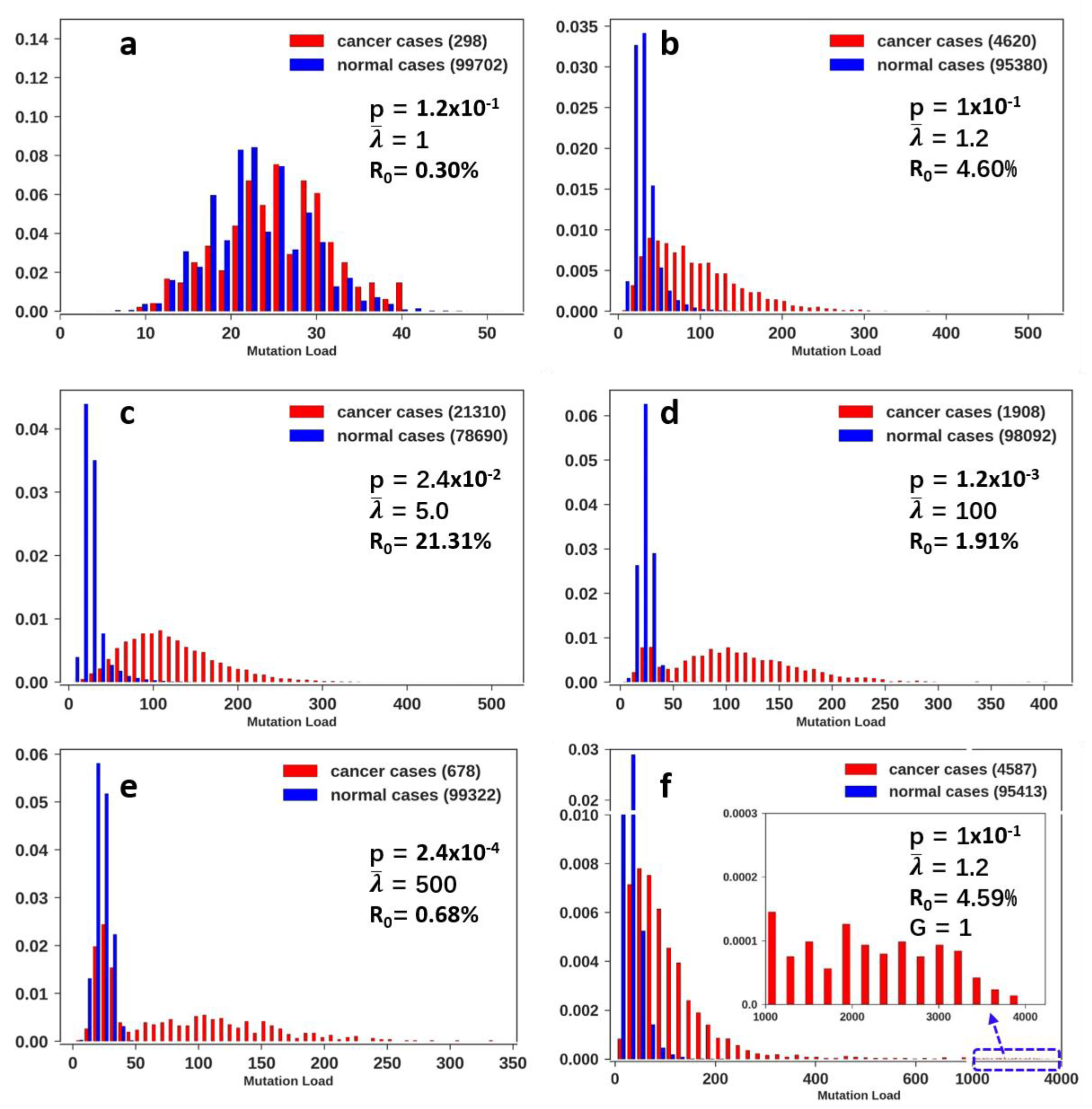
Mutation load distribution across mutator effect sizes. (a-e) Normalized histogram of mutation load with 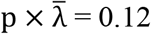 where p is the probability of mutator mutations and 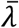 is the mean mutator strength. a – e: the mutator strength continues to increase but the mutators become less prevalent (see the main text). R_0_ represents the lifetime cancer risk, i.e. the probability of becoming cancerous by accumulating at least five cancer drivers. For each parameter set, we simulate 100000 cases. (f) The same as panel (b) except for an additional parameter of G=1 (year), which is the latency time between the acquisition of the five driver mutations and the exponential expansion of the cancer cell population. Recent evidence has increasingly suggested the existence of this latency time due to intense local competitions among cell clones (Martincorena, et al. 2018; Yizhak, et al. 2019; Zhu, et al. 2019). During this time gap, mutation accumulation may also accelerate, thus giving rise to the strong right-skewed distribution.

In Fig. 3, the lifetime cancer risk (R_0_) and mutation load both increase as the mutator strength increases (Fig. 3a-c). However, above a certain level, the risk starts to decrease (Fig. 3c-e). This reversal is due to the rareness of strong mutators, when most cases do not acquire even a single mutator. Overall, many weak mutators are much more compatible with the observed distributions of mutation load than a few strong ones (Fig. 3a-c and Fig. 3d-e vs. Fig. 1a).

While the model predicts a long tail of high mutation load, the maximal number of mutations is still smaller than 500 (Fig. 3b-e) whereas the observations of Fig. 1a show a much longer and thinner tail. In the framework of the runaway model, a very long tail is possible, provided that there is time for mutations to accrue. For example, there may exist a time gap (G, also known as incubation period or latency time (Nadler and Zurbenko 2014)) between the emergence of the single progenitor cancer cell and its subsequent expansion. Even with a small time gap (G = 1 year), the maximum number of mutations in tumors can reach nearly 4000 (Fig. 3f). The distribution with G =1 indeed agrees well with the observations on colorectal cancer (Fig. 3f vs. Fig. 1a).

### IV. Evolution of the germline mutation rate (*μ*_0_)

We now transition from analyzing somatic mutations to predicting the germline mutation rate. Unlike in the soma, all germline mutations in the gamete are present in every cell of the progeny. For this reason, deleterious mutations in the germline would have a much greater fitness consequence than somatic mutations and the germline mutation rate is expected to be lower than in the soma. The question is “how much lower”. It is then natural to ask how the baseline mutation rate, *μ*_0_, would affect the runaway process in the germline. After all, a higher *μ*_0_ is equivalent to the presence of some mutators.

This rate of germline mutation in humans, referred to as *μ*\*, is estimated to be ~ 1 *de novo* mutations per year per gamete, corresponding to about 0.4 × 10^−9^ /bp /year (Scally and Durbin 2012; Jonsson, et al. 2017). In this section, we are interested in the coding regions and set *μ*\* = 0.02 per year. We then ask about the consequence of mutation accumulation if *μ*_0_ is larger than *μ*\*, with *μ*_0_ = 2*μ*\*, 5*μ*\* or 10*μ*\*, in comparison with *μ*_0_ = *μ*\*. The average strength of the mutator, 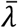, is assumed to be either 2 or 5. Given that *μ*\* = 0.4 × 10^−9^/bp/year is a very small number to begin with, a 2 or 5 fold increase would still yield a very small mutation rate.

The probability of runaway accumulation will be referred to as the “fitness risk”, which is the average fitness reduction in the progeny (see Fig. 4). Given the effective population size of humans in the range of (5-10) × 10^3^, a fitness reduction of 0.005 - 0.01 should be considered medium to high. Indeed, the calculated fitness risk due to deleterious mutations is ~ 0.5 × 10^−4^, based on human polymorphism data (see Table 1 and Methods). Therefore, the fitness risk in extant humans is minimal if the baseline mutation rate does not evolve.

**Table 1.**
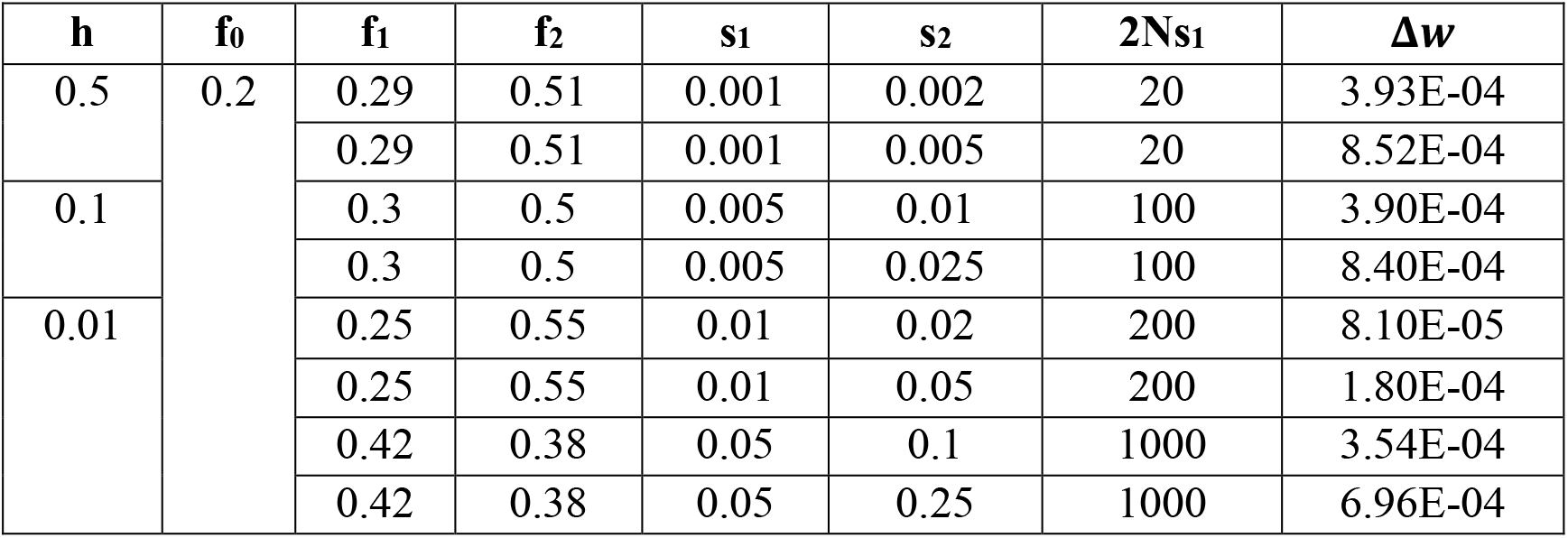
Estimation of fitness risk of human population

**Fig. 4.**
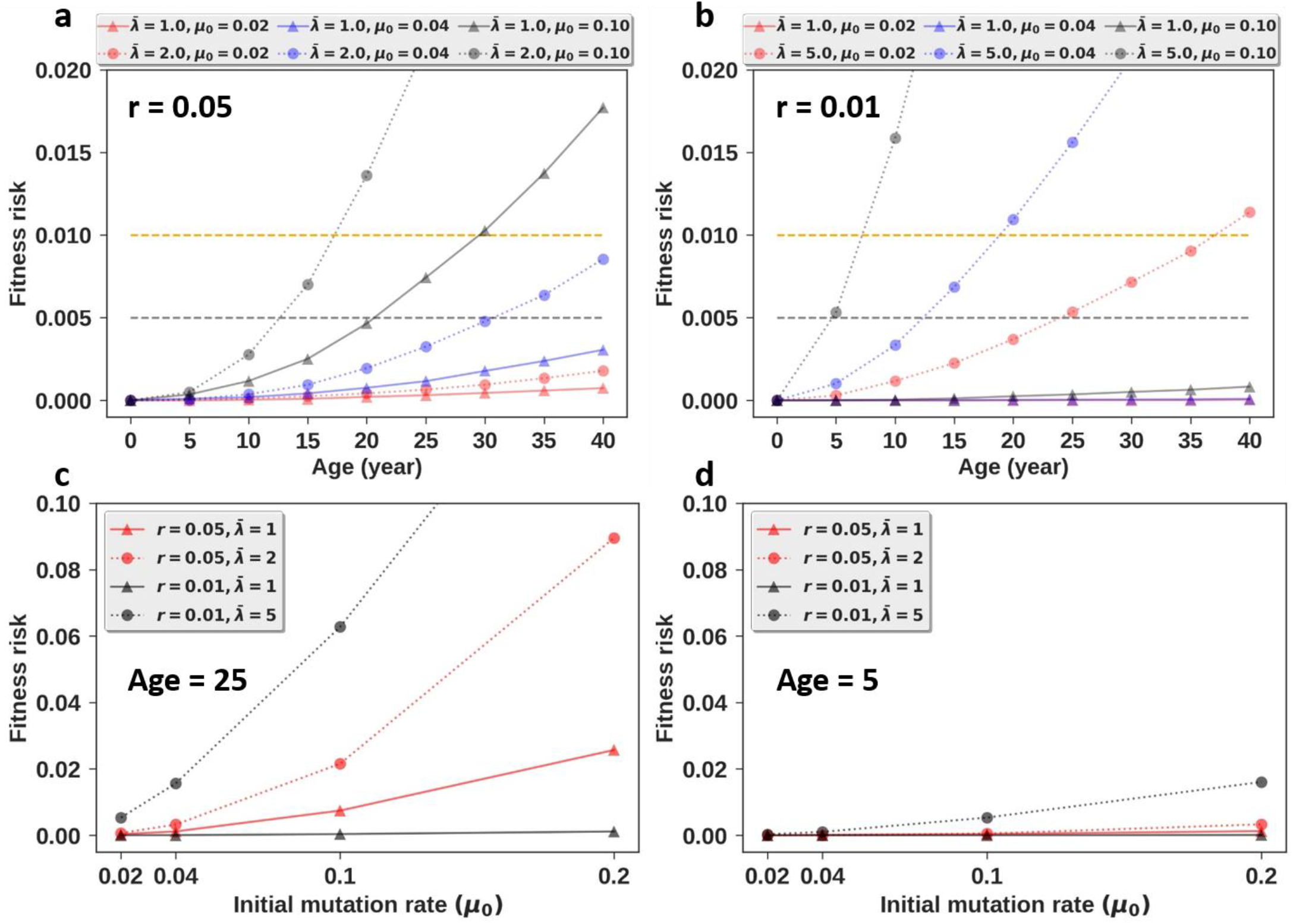
Probability of fitness reduction (or risk) as a function of the germline mutation rate at age 0, *μ*_0_. This value in extant humans is estimated to be 0.02. We use *μ*_0_= 0.02, 0.04, 0.10, 0.20 in the simulations. p = 0.1 for all panels. (a) Fitness reduction due to runaway mutation accumulation under frequent fitness-altering mutations (r = 0.05) and weak mutators 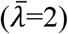. (b) As in (a), but with less frequent fitness-altering variants (r = 0.01) and stronger mutators 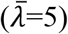. (c, d) Fitness reduction at age 25 or 5 as a function of the baseline mutation rate. The low fitness values in these panels are given numerically in Table S1, S2.

Results from two schemes of mutator effect are presented. In scheme A with 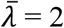 and r = 0.05, fitness-altering mutations are common but mutator strength is moderate (Fig. 4a; Table S1). In this scheme and in the absence of mutators, the risk stays low until age 40 if *μ*_0_ = 2*μ*\*. Nevertheless, because the fitness mutations are common, the risk would be moderate at age 25 if *μ*_0_ = 5*μ*\*. With mutators, if *μ*_0_ = 2*μ*\*, the risk would be moderate at age 30 and high at age 40. If *μ*_0_ = 5*μ*\*, then the risk is already high at age 18.

In scheme B with 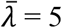 and r = 0.01, the fitness-altering mutations are less common but the mutators are strong (Fig. 4b; Table S2). We shall now consider the fitness risk if there are no mutators. First, according to simulations, the fitness risk would be imperceptible for all parameter values (Fig. 4b; Table S2). Second, when we use the actual polymorphism data on deleterious mutations, the calculated fitness risk would remain < 10^−3^ even if *μ*_0_ = 10*μ*\* (see Methods). Hence, the baseline mutation rate in humans appears to be far below what natural selection would tolerate, if there are no mutators. In contrast, if we consider mutators with the associated risk of runaway process, even *μ*_0_ = *μ*\* would incur a moderate risk of 0.005 at age 25. If *μ*_0_ = 2*μ*\*, the risk is moderate at age 13 (probably before sexual maturity) and high at age 19. With strong mutators and *μ*_0_ = 5*μ*\*, the fitness risk would be exceedingly high at age 10.

The trend is most discernible if we focus on the risk at age 25, assumed to be the mean reproductive age of humans (Fig. 4c). With mutators (the top two lines), a two-fold increase in the baseline mutation rate would make the risk rather high at age 25. A five-fold increase to *μ*_0_ = 0.1 would make the risk of fitness reduction unacceptable. On the other hand, without the runaway process, the baseline mutation could be two, or even five-fold, higher than its current level, regardless of the proportion of lethal mutations. The lowest line shows that, given r = 0.01, *μ*_0_ can easily be 10-fold higher than it is now if there is no threat of runaway mutation. In conclusion, mutators could set a limit on the baseline mutation rate, *μ*_0_, that may otherwise seem too low.

The threat of runaway accumulation is much reduced if reproduction happens much earlier. For example, at age five, the cost of letting *μ*_0_ increase by even 10-fold would be modest (Fig. 4d). This age-dependency could explain why the mutation rate per year in mice is 10 times higher than in humans (Lindsay, et al. 2018). Furthermore, there is a negative correlation between generation time and per year mutation rate. Comparing Fig. 4d with Fig. 4c, one may make a case that the reduced risk of runaway mutation accumulation may permit the baseline mutation rate to rise high in taxa with shorter generation time. In contrast, long-living species like humans may have set *μ*_0_ low to avoid mutation rates increasing at an accelerating pace. Indeed, in more than 1500 trios in Iceland where the parental age rarely exceeds 50, no runaway accumulation was observed (Jonsson, et al. 2017).

## Discussion

Mutation is unique among evolutionary forces as mutations themselves form a positive feedback loop. With mutator mutations begetting more mutations, the process can evolve very quickly, even within a lifetime. Indeed, somatic tissues with a high mutation load often evolve into tumors (Loeb 1991; Hanahan and Weinberg 2011; Cancer Genome Atlas 2012; Cancer Genome Atlas Research, et al. 2013). The source of mutators is crucial for the runaway process. An interesting example is a panel of 66 DNA polymerase I mutants in *E. coli* that yield replication fidelities spanning three orders of magnitude (Loh, et al. 2010). Such strong mutators, however, are not found to drive high mutation rates as shown in Fig. 1, thus suggesting the source of mutator mutations to be rather diverse.

Until now, mutators driving a runaway process have not yet been documented for germline evolution. We thus re-evaluate the assumption of an unvarying mutation rate across time. In the evolutionary time span, the postulate of a constant mutation rate driving a molecular clock has been rejected (Wu and Li 1985; Kumar and Subramanian 2002; Kumar 2005; Elango, et al. 2006; Moorjani, et al. 2016). Furthermore, by tracking the change of the α ratio through time (α being the mutation rate in males vs. that in females), Makova and Li report the α ratio in the extant hominids to be much smaller than in the earlier period of primate evolution (Makova and Li 2002). More recently, the mutation rate in extant humans, observed by the parents-offspring trio sequencing, is found to be only 1/3 as large as those of other apes – chimpanzee, gorilla, and orangutan (Besenbacher, et al. 2019). These observations suggest that the mutation rate has been evolving rapidly.

Since the mutation rate evolves, the extant rate should reflect past selective pressure (Lynch 2010, 2011; Scally and Durbin 2012; Sung, et al. 2012; Wielgoss, et al. 2013). However, the germline mutation rate in extant humans seems unnecessarily low (Fig. 4). The apparent excess in replication fidelity can be resolved if the avoidance of runaway mutation accumulation is factored in. As the potential for such accumulation should increase with age, one may ask for evidence for this process in men of advanced age, say, > 70. There are hints that the accumulation of mutations is closer to exponential than linear when very old paternal parents are included (Risch, et al. 1987; Wyrobek, et al. 2006; Kong, et al. 2012; Khandwala, et al. 2018). In this context, the issue of advanced paternal age would have theoretical, medical, and social-cultural significance.

The avoidance of runaway mutation accumulation may also explain why the average age of reproduction has such an impact on the baseline mutation rate. In general, short-living animals have a much higher baseline rate (Welch, et al. 2008; Thomas, et al. 2010; Wilson Sayres, et al. 2011). For example, the per year mutation rate in mice is 10 times higher than in humans (Lindsay, et al. 2018). Even among hominoids, the slight delay in the age of reproduction (~ age 29) may have contributed to the much lower baseline rate in humans than in other apes (age 19-25) (Langergraber, et al. 2012). In Fig. 3f, it is shown that one extra year can extend the tail length many-fold. As pointed out (Besenbacher, et al. 2019), this strong generation-time dependence may explain why the mutation rate obtained by trio sequencing in humans is only half as large as that inferred from the evolutionary analysis between human and other apes. Given their longer generation time, modern humans appear to have a lower baseline mutation rate than their ancestors and relatives.

In conclusion, the evolution of the mutation rate could be a highly dynamic process due to the positive feedback of mutators on mutations. This mechanism is revealed by cancer genomic studies (Cancer Genome Atlas 2012; Cancer Genome Atlas Research, et al. 2013; Kandoth, et al. 2013; Lawrence, et al. 2014). The process of mutation is another example, in addition to genetic drift (Chen, et al. 2017), natural selection (Chen, et al. 2019), and sex chromosome evolution (Xu, et al. 2017) that cancer evolution studies can enrich the general theories of biological evolution (Wu, et al. 2016; Wen, et al. 2018).

## Methods

### I. Data Handling

#### Estimating the proportions of DNA repair genes and cancer driver genes in the genome

In the runaway model, *p* is the proportion of mutator mutations and *r* is the proportion of fitness-altering mutations. Both have to be estimated. We first use the proportion of DNA repair genes as a lower bound on *p*. For fitness mutations, we use the cancer driver mutations compiled from the literature.

The 177 DNA repair genes and 299 cancer driver genes were obtained from the public databases as well as published papers (Wood, et al. 2001; Wood, et al. 2005; Lawrence, et al. 2014; Bailey, et al. 2018). We use the software GTFtools (Li 2018) to calculate non-overlapping exon length for these genes (see Table S3, Table S4). The total lengths of exons of the 177 DNA repair genes and the 299 cancer driver genes are 1026236 bp and 2372955 bp, respectively. The exome of the human genome consists of roughly 180,000 exons constituting about 1% of the total genome (Ng, et al. 2009). Thus, the exome proportion of DNA repair genes is 1026326 / (3.2×10^7^) = 0.032 and 2372955 / (3.2×10^7^) = 0.074 for cancer driver genes. Considering that the ratio of nonsynonymous mutations over synonymous mutations is around 2:1, the proportion of function-changing sites in DNA repair genes is 0.02 and 0.04 in cancer driver genes.

#### Extraction of SNVs from whole exome (or genome) data from cancerous and normal tissues

We obtained exome mutation data called by Mutect2 and the associated clinical information from TCGA. There are 10429 patients across 33 cancer types. For each case, we only care about single nucleotide variants (SNVs) in exons and discard all other mutations. We divide the SNVs into synonymous (”5’UTR”, “3’UTR”, “Silent”) and nonsynonymous mutations (”Missense Mutation”, “Nonsense Mutation”, “Nonstop Mutation”, “Translation Start Site”, “Splice Site”, “Splice Region”). We then counted the numbers of nonsynonymous and synonymous mutations in the 177 DNA repair and 299 cancer driver genes in each patient. To compare with the contribution of DNA repair mutations to the mutation load, we randomly chose 207 genes with the same length as the 177 DNA repair genes and likewise counted the nonsynonymous and synonymous mutations.

We obtained filtered vcf files (whole genome sequencing data) corresponding to 45 normal tissue samples (21 colons, 14 small intestines, and 10 livers) from Blokzijl et al. (Blokzijl, et al. 2016). We used ANNOVAR (Yang and Wang 2015) for variant annotation. We processed these data similarly to the TCGA data to obtain nonsynonymous and synonymous mutation counts in DNA repair, cancer driver, and random genes.

### II. Model and simulation

The structure of the runaway mutation accumulation model is presented in the main text and the approximate analytical solutions are given in the Supplemental text. We describe the simulation procedures below.

#### Simulation steps

1. Initialization. Determine the lifetime (*T*) of an individual. Set initial age t = 0, initial mutation rate *μ* = *μ*_0_, the number of mutator mutations *n*_*k*_ = 0, the number of mutations 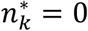, the number of fitness-altering mutations K = 0.
2. Determine the waiting time Δt to a new mutation by sampling from the exponential distribution with mean 1/*μ*.
3. The new mutations are mutators with probability p, and fitness-altering with probability r. p and r are mutually exclusive. If the new mutation is fitness-altering, go to step 4. If the new mutation is a mutator, go to step 5.
4. Fitness mutation: Update fitness-altering mutation number K = K + 1.
5. Mutator mutation: Sample a value k from the exponential distribution with mean 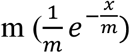 (see Mutator strength below). Update mutation rate *μ* = *μ* × *e*^*k*^. Update mutator mutation number *n*_*k*_ = *n*_*k*_ + 1.
6. Update t = t + Δt, 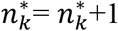. If we are simulating the mutation accumulation in somatic tissues (i.e. the tumorigenesis process), go to step 7. If in the germline, go to step 8.
7. According to the tumor onset function (see main text), determine whether the individual forms a tumor. If a tumor is formed or *t* > *T*, stop simulation (Note: if there is a latency time G, the cancer case can also accumulate mutations during latency). Otherwise, return to step 2.
8. If K > 1 (synthetic lethal) or *t* > *T*, stop simulation. Otherwise, return to step 2.

For each parameter space, we simulate 100000 cases unless otherwise specified.

#### Mutator strength

In the main text, the effect of a new mutator on the mutation rate is

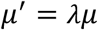

We assume that *x* = *ln*(*λ*) follows an exponential distribution with mean *m*. *λ* thus follows the log-exponential distribution. A good estimation for the mutator strength is the mean *λ*. Here we just obtain the mean of the log-exponential variable (*λ*) to estimate the average effect of a new mutator.

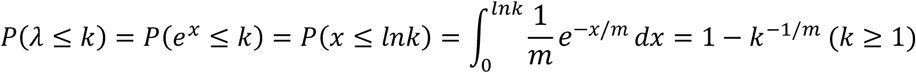

i.e. the cumulative density function (CDF) of *λ* is

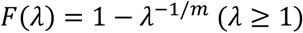

Taking the derivative gives the density of *λ*,

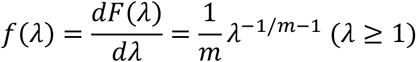

Then the mean of *λ* is given by

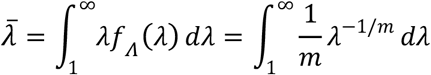

When 0 < *m* < 1,

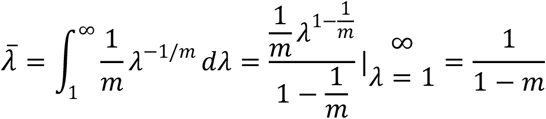

When *m* = 1,

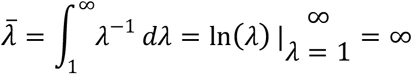

When *m* > 1,

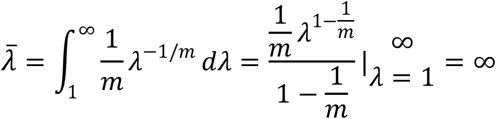

The mean of *λ* goes to infinity when *m* ≥ 1. However, the mutator effect, *λ*, of a new mutator cannot be infinite in our case. In addition, when m is small,

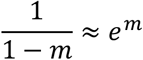

Thus, we just let the mean effect of a mutator approximately be

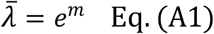

### III. Estimation of fitness risk (or reduction) from human polymorphism data

Here we calculate the fitness reduction of the human population according to the proportion and selective coefficients of deleterious mutations estimated by Fay et al (Fay, et al. 2001). Like Fay et al, we assume the proportions of neutral, slightly deleterious, and strongly deleterious mutations to be f_0_, f_1_, and f_2_ respectively. The average selection coefficient of slightly deleterious mutations is s_1_ and s_2_ for strongly deleterious variants. We denote the dominance coefficient by h, mutation rate per year by *μ*_0_, and the conception time by T. We let effective population size of humans be N = 10000. Then the average number of new mutations of an individual after a single generation is

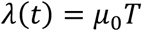

Assuming that the mutation load follows the Poisson distribution, the probability of an individual with *i* new mutations is given by

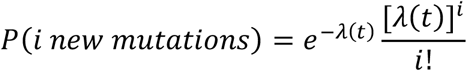

In the infinite allele model, two mutations are impossible at the same allele. Thus, the new mutation will be in a heterozygous state and the expected fitness (multiplicative epistasis) of an individual with the new mutation *i* is

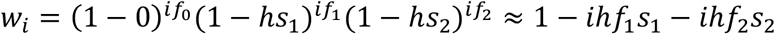

Further, the average fitness of the population is

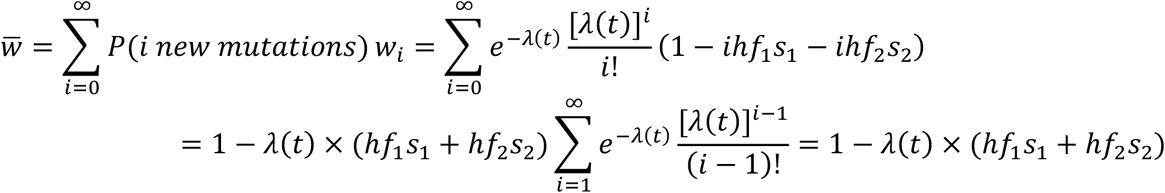

Thus, the fitness reduction of the human population can be obtained by

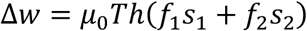

Comparing with the human polymorphism data, Fay et al. used the deleterious model to estimate the following parameters: f_0_ ~ 0.2, f_1_~0.3, f_2_~0.5, 2Nhs_1_~10 (Fay, et al. 2001). We use several values for these parameters close to the estimated numbers and assume s_2_ is two or five-fold higher than s_1_. We hen let *μ*_0_=0.02 and *T*=30 and use the preceding equation to estimate the corresponding fitness reduction (Table 1). The table shows that fitness reduction due to the baseline mutation rate (*μ*_0_=0.02) will be ~ 0.5 × 10^−4^. According to the preceding equation, the fitness reduction will remain < 10^−3^ (0.5 × 10^−3^) even if we increase the mutation rate 10-fold. Thus, the estimated mutation rate (~0.02 per exome per year) in humans appears to be far below what natural selection would tolerate.

## Acknowledgements

We would like to thank Xionglei He, Zheng Hu, and the members of Wu Lab for discussions and advices. This work was supported by the National Natural Science Foundation of China (31730046 and 91731301) and the 985 Project (33000-18841204).

## References

Alexandrov LB, Jones PH, Wedge DC, Sale JE, Campbell PJ, Nik-Zainal S, Stratton MR. 2015. Clock-like mutational processes in human somatic cells. Nat Genet 47:1402–1407.

Bae T, Tomasini L, Mariani J, Zhou B, Roychowdhury T, Franjic D, Pletikos M, Pattni R, Chen BJ, Venturini E, et al. 2018. Different mutational rates and mechanisms in human cells at pregastrulation and neurogenesis. Science 359:550–555.

Bailey MH, Tokheim C, Porta-Pardo E, Sengupta S, Bertrand D, Weerasinghe A, Colaprico A, Wendl MC, Kim J, Reardon B, et al. 2018. Comprehensive Characterization of Cancer Driver Genes and Mutations. Cell 173:371–385 e318.

Besenbacher S, Hvilsom C, Marques-Bonet T, Mailund T, Schierup MH. 2019. Direct estimation of mutations in great apes reconciles phylogenetic dating. Nat Ecol Evol 3:286–292.

Blokzijl F, de Ligt J, Jager M, Sasselli V, Roerink S, Sasaki N, Huch M, Boymans S, Kuijk E, Prins P, et al. 2016. Tissue-specific mutation accumulation in human adult stem cells during life. Nature 538:260–264.

Cancer Genome Atlas N. 2012. Comprehensive molecular characterization of human colon and rectal cancer. Nature 487:330–337.

Cancer Genome Atlas Research N, Weinstein JN, Collisson EA, Mills GB, Shaw KR, Ozenberger BA, Ellrott K, Shmulevich I, Sander C, Stuart JM. 2013. The Cancer Genome Atlas Pan-Cancer analysis project. Nat Genet 45:1113–1120.

Charlesworth D, Charlesworth B, Marais G. 2005. Steps in the evolution of heteromorphic sex chromosomes. Heredity (Edinb) 95:118–128.

Chen B, Shi Z, Chen Q, Shen X, Shibata D, Wen H, Wu CI. 2019. Tumorigenesis as the Paradigm of Quasineutral Molecular Evolution. Mol Biol Evol 36:1430–1441.

Chen Y, Tong D, Wu CI. 2017. A New Formulation of Random Genetic Drift and Its Application to the Evolution of Cell Populations. Mol Biol Evol 34:2057–2064.

Drake JW. 1991. A constant rate of spontaneous mutation in DNA-based microbes. Proc Natl Acad Sci U S A 88:7160–7164.

El Meouche I, Dunlop MJ. 2018. Heterogeneity in efflux pump expression predisposes antibiotic-resistant cells to mutation. Science 362:686–690.

Elango N, Thomas JW, Program NCS, Yi SV. 2006. Variable molecular clocks in hominoids. Proc Natl Acad Sci U S A 103:1370–1375.

Eyre-Walker A, Keightley PD. 2007. The distribution of fitness effects of new mutations. Nat Rev Genet 8:610–618.

Eyre-Walker A, Woolfit M, Phelps T. 2006. The distribution of fitness effects of new deleterious amino acid mutations in humans. Genetics 173:891–900.

Fay JC, Wyckoff GJ, Wu CI. 2001. Positive and negative selection on the human genome. Genetics 158:1227–1234.

Fearon ER, Vogelstein B. 1990. A genetic model for colorectal tumorigenesis. Cell 61:759–767.

Hanahan D, Weinberg RA. 2011. Hallmarks of cancer: the next generation. Cell 144:646–674.

Jonsson H, Sulem P, Kehr B, Kristmundsdottir S, Zink F, Hjartarson E, Hardarson MT, Hjorleifsson KE, Eggertsson HP, Gudjonsson SA, et al. 2017. Parental influence on human germline de novo mutations in 1,548 trios from Iceland. Nature 549:519–522.

Ju YS, Martincorena I, Gerstung M, Petljak M, Alexandrov LB, Rahbari R, Wedge DC, Davies HR, Ramakrishna M, Fullam A, et al. 2017. Somatic mutations reveal asymmetric cellular dynamics in the early human embryo. Nature 543:714–718.

Kandoth C, McLellan MD, Vandin F, Ye K, Niu B, Lu C, Xie M, Zhang Q, McMichael JF, Wyczalkowski MA, et al. 2013. Mutational landscape and significance across 12 major cancer types. Nature 502:333–339.

Khandwala YS, Baker VL, Shaw GM, Stevenson DK, Lu Y, Eisenberg ML. 2018. Association of paternal age with perinatal outcomes between 2007 and 2016 in the United States: population based cohort study. BMJ 363:k4372.

Kimura M. 1983. The neutral theory of molecular evolution. Cambridge: Cambridge University Press.

Kong A, Frigge ML, Masson G, Besenbacher S, Sulem P, Magnusson G, Gudjonsson SA, Sigurdsson A, Jonasdottir A, Jonasdottir A, et al. 2012. Rate of de novo mutations and the importance of father’s age to disease risk. Nature 488:471–475.

Kumar S. 2005. Molecular clocks: four decades of evolution. Nat Rev Genet 6:654–662.

Kumar S, Subramanian S. 2002. Mutation rates in mammalian genomes. Proc Natl Acad Sci U S A 99:803–808.

Langergraber KE, Prufer K, Rowney C, Boesch C, Crockford C, Fawcett K, Inoue E, Inoue-Muruyama M, Mitani JC, Muller MN, et al. 2012. Generation times in wild chimpanzees and gorillas suggest earlier divergence times in great ape and human evolution. Proc Natl Acad Sci U S A 109:15716–15721.

Lawrence MS, Stojanov P, Mermel CH, Robinson JT, Garraway LA, Golub TR, Meyerson M, Gabriel SB, Lander ES, Getz G. 2014. Discovery and saturation analysis of cancer genes across 21 tumour types. Nature 505:495–501.

Lawrence MS, Stojanov P, Polak P, Kryukov GV, Cibulskis K, Sivachenko A, Carter SL, Stewart C, Mermel CH, Roberts SA, et al. 2013. Mutational heterogeneity in cancer and the search for new cancer-associated genes. Nature 499:214–218.

Li H. 2018. GTFtools: a Python package for analyzing various modes of gene models. bioRxiv.

Li W-H. 2007. Molecular evolution. New York; Basingstoke: W.H. Freeman; Palgrave [distributor].

Li WH, Wu CI, Luo CC. 1985. A new method for estimating synonymous and nonsynonymous rates of nucleotide substitution considering the relative likelihood of nucleotide and codon changes. Mol Biol Evol 2:150–174.

Lindsay SJ, Rahbari R, Kaplanis J, Keane T, Hurles M. 2018. Striking differences in patterns of germline mutation between mice and humans. bioRxiv:082297.

Lodato MA, Rodin RE, Bohrson CL, Coulter ME, Barton AR, Kwon M, Sherman MA, Vitzthum CM, Luquette LJ, Yandava CN, et al. 2018. Aging and neurodegeneration are associated with increased mutations in single human neurons. Science 359:555–559.

Loeb LA. 1991. Mutator Phenotype May Be Required for Multistage Carcinogenesis. Cancer Res 51:3075–3079.

Loh E, Salk JJ, Loeb LA. 2010. Optimization of DNA polymerase mutation rates during bacterial evolution. Proc Natl Acad Sci U S A 107:1154–1159.

Lynch M. 2010. Evolution of the mutation rate. Trends Genet 26:345–352.

Lynch M. 2011. The lower bound to the evolution of mutation rates. Genome Biology and Evolution 3:1107–1118.

Lynch M, Ackerman MS, Gout JF, Long H, Sung W, Thomas WK, Foster PL. 2016. Genetic drift, selection and the evolution of the mutation rate. Nat Rev Genet 17:704–714.

Makova KD, Li WH. 2002. Strong male-driven evolution of DNA sequences in humans and apes. Nature 416:624–626.

Martincorena I, Fowler JC, Wabik A, Lawson ARJ, Abascal F, Hall MWJ, Cagan A, Murai K, Mahbubani K, Stratton MR, et al. 2018. Somatic mutant clones colonize the human esophagus with age. Science 362:911–917.

Martincorena I, Raine KM, Gerstung M, Dawson KJ, Haase K, Van Loo P, Davies H, Stratton MR, Campbell PJ. 2017. Universal Patterns of Selection in Cancer and Somatic Tissues. Cell 171:1029–1041 e1021.

Martincorena I, Roshan A, Gerstung M, Ellis P, Van Loo P, McLaren S, Wedge DC, Fullam A, Alexandrov LB, Tubio JM, et al. 2015. Tumor evolution. High burden and pervasive positive selection of somatic mutations in normal human skin. Science 348:880–886.

Milholland B, Auton A, Suh Y, Vijg J. 2015. Age-related somatic mutations in the cancer genome. Oncotarget 6:24627–24635.

Moorjani P, Amorim CE, Arndt PF, Przeworski M. 2016. Variation in the molecular clock of primates. Proc Natl Acad Sci U S A 113:10607–10612.

Nadler DL, Zurbenko IG. 2014. Estimating Cancer Latency Times Using a Weibull Model. Advances in Epidemiology Advances in Epidemiology 2014:1–8.

Ng SB, Turner EH, Robertson PD, Flygare SD, Bigham AW, Lee C, Shaffer T, Wong M, Bhattacharjee A, Eichler EE, et al. 2009. Targeted capture and massively parallel sequencing of 12 human exomes. Nature 461:272–276.

Podolskiy DI, Lobanov AV, Kryukov GV, Gladyshev VN. 2016. Analysis of cancer genomes reveals basic features of human aging and its role in cancer development. Nat Commun 7:12157.

Risch N, Reich EW, Wishnick MM, McCarthy JG. 1987. Spontaneous mutation and parental age in humans. Am J Hum Genet 41:218–248.

Scally A, Durbin R. 2012. Revising the human mutation rate: implications for understanding human evolution. Nat Rev Genet 13:745–753.

Schuster-Bockler B, Lehner B. 2012. Chromatin organization is a major influence on regional mutation rates in human cancer cells. Nature 488:504–507.

Segurel L, Wyman MJ, Przeworski M. 2014. Determinants of mutation rate variation in the human germline. Annu Rev Genomics Hum Genet 15:47–70.

Sniegowski PD, Gerrish PJ, Lenski RE. 1997. Evolution of high mutation rates in experimental populations of E. coli. Nature 387:703–705.

Sung W, Ackerman MS, Miller SF, Doak TG, Lynch M. 2012. Drift-barrier hypothesis and mutation-rate evolution. Proc Natl Acad Sci U S A 109:18488–18492.

Thomas GWC, Wang RJ, Puri A, Harris RA, Raveendran M, Hughes DST, Murali SC, Williams LE, Doddapaneni H, Muzny DM, et al. 2018. Reproductive Longevity Predicts Mutation Rates in Primates. Current Biology 28:3193–3197 e3195.

Thomas JA, Welch JJ, Lanfear R, Bromham L. 2010. A generation time effect on the rate of molecular evolution in invertebrates. Mol Biol Evol 27:1173–1180.

Welch JJ, Bininda-Emonds OR, Bromham L. 2008. Correlates of substitution rate variation in mammalian protein-coding sequences. BMC Evol Biol 8:53.

Wen HJ, Wang HY, He XL, Wu CI. 2018. On the low reproducibility of cancer studies. National Science Review 5:619–624.

Wielgoss S, Barrick JE, Tenaillon O, Wiser MJ, Dittmar WJ, Cruveiller S, Chane-Woon-Ming B, Medigue C, Lenski RE, Schneider D. 2013. Mutation rate dynamics in a bacterial population reflect tension between adaptation and genetic load. Proc Natl Acad Sci U S A 110:222–227.

Wilson Sayres MA, Venditti C, Pagel M, Makova KD. 2011. Do variations in substitution rates and male mutation bias correlate with life-history traits? A study of 32 mammalian genomes. Evolution 65:2800–2815.

Wood RD, Mitchell M, Lindahl T. 2005. Human DNA repair genes, 2005. Mutat Res 577:275–283.

Wood RD, Mitchell M, Sgouros J, Lindahl T. 2001. Human DNA repair genes. Science 291:1284–1289.

Wu CI, Li WH. 1985. Evidence for higher rates of nucleotide substitution in rodents than in man. Proc Natl Acad Sci U S A 82:1741–1745.

Wu CI, Wang HY, Ling S, Lu X. 2016. The Ecology and Evolution of Cancer: The Ultra-Microevolutionary Process. Annu Rev Genet 50:347–369.

Wyrobek AJ, Eskenazi B, Young S, Arnheim N, Tiemann-Boege I, Jabs EW, Glaser RL, Pearson FS, Evenson D. 2006. Advancing age has differential effects on DNA damage, chromatin integrity, gene mutations, and aneuploidies in sperm. Proc Natl Acad Sci U S A 103:9601–9606.

Xu J, Peng X, Chen Y, Zhang Y, Ma Q, Liang L, Carter AC, Lu X, Wu CI. 2017. Free-living human cells reconfigure their chromosomes in the evolution back to uni-cellularity. Elife 6:e28070.

Yang H, Wang K. 2015. Genomic variant annotation and prioritization with ANNOVAR and wANNOVAR. Nature Protocols 10:1556–1566.

Yizhak K, Aguet F, Kim J, Hess JM, Kubler K, Grimsby J, Frazer R, Zhang H, Haradhvala NJ, Rosebrock D, et al. 2019. RNA sequence analysis reveals macroscopic somatic clonal expansion across normal tissues. Science 364:eaaw0726.

Zapata L, Pich O, Serrano L, Kondrashov FA, Ossowski S, Schaefer MH. 2018. Negative selection in tumor genome evolution acts on essential cellular functions and the immunopeptidome. Genome Biology 19:67.

Zhang L, Dong X, Lee M, Maslov AY, Wang T, Vijg J. 2019. Single-cell whole-genome sequencing reveals the functional landscape of somatic mutations in B lymphocytes across the human lifespan. Proc Natl Acad Sci U S A 116:9014–9019.

Zhu M, Lu T, Jia Y, Luo X, Gopal P, Li L, Odewole M, Renteria V, Singal AG, Jang Y, et al. 2019. Somatic Mutations Increase Hepatic Clonal Fitness and Regeneration in Chronic Liver Disease. Cell 177:608–621 e612.

