## supplementary information for "Mutations beget more mutations – The baseline mutation rate and runaway accumulation"

**Supplementary Information includes:**

Approximate analytical solutions for runaway model

Fig. S1 to S4

Table S1 to S4

References for SI references citations

**Approximate analytical solutions for runaway model**

Here we obtain the analytical solutions for the runaway model. The runaway model takes the general form:

$$P\left( n_{k}^{*}(t+\Delta t)=x|n_{k}^{*}\left( t \right) \right)=e^{-\mu\left( t \right)\Delta t}\frac{{(\mu\left( t \right)\Delta t)}^{x-n_{k}^{*}\left( t \right)}}{\left( x-n_{k}^{*}\left( t \right) \right)!}$$

where $n_{k}^{*}\left( t \right)$ is the number of total mutations at age t. To solve the above equation, the key step is to get the mutation rate $\mu\left( t \right)$. After a small time interval $\Delta t$, the average number of new mutator mutations is

$$\Delta n_{k}=n_{k}\left( t+\Delta t \right)-n_{k}\left( t \right)=\mu(t)\Delta t\times p$$

where $n_{k}\left( t \right)$ is the number of mutator variants at age t. According to $Eq (A1)$ in Methods, the corresponding mutator effect of the new mutator mutations is approximately equal to

$$\bar{\lambda}\approx e^{m\Delta n_{k}}=e^{pm\mu(t)\Delta t}$$

So the mutation rate changes to

$$\mu(t+\Delta t)=\bar{\lambda}\mu(t)=\mu(t)e^{pm\mu(t)\Delta t}$$

After transformation, we get

$$\frac{\mu(t+\Delta t)-\mu(t)}{\Delta t}=\frac{\mu(t)(e^{pm\mu(t)\Delta t}-1)}{\Delta t}$$

Letting $\Delta t\to0$, we get the following differential equation

$$\frac{d\mu(t)}{dt}=pm{\mu(t)}^{2}$$

Then the average mutation rate across time is:

$$\mu\left( t \right)=\frac{\mu_{0}}{1-{mp\mu}_{0}t} Eq. (A2)$$

It is important to emphasize that the range of age t is $0\leq t<\frac{1}{mp\mu_{0}}$.

$$\lim_{t\to\frac{1}{mp\mu_{0}}} \frac{\mu_{0}}{1-{mp\mu}_{0}t}=\infty$$

That is to say, the average mutation rate will be very large when the age of an individual reaches $\frac{1}{mp\mu_{0}}$.

Now the number of mutations follows a nonhomogeneous Poisson process with variate rate $\mu\left( t \right)$. According to the properties of this process (Grimmett and Stirzaker 2009), the number of mutations at age t also follows a Poisson process, whose rate is the average number of mutations at age t:

$$\lambda\left( t \right)=\int_{0}^{t} \mu\left( x \right)dx=-\frac{ln\left( 1-{mp\mu}_{0}t \right)}{mp} Eq. (A3)$$

Then the probability that an individual has $x$ mutations at age t is

$$P\left( n_{k}^{*}\left( t \right)=x \right)=e^{-\lambda\left( t \right)}\frac{{\lambda\left( t \right)}^{x}}{x!}=e^{\frac{ln\left( 1-{mp\mu}_{0}t \right)}{mp}}\frac{{(-\frac{ln\left( 1-{mp\mu}_{0}t \right)}{mp})}^{x}}{x!} Eq. (A4.1)$$

Similarly, the probability density function of mutator mutations number $n_{k}(t)$ and fitness-altering mutations number $K(t)$:

$$P\left( n_{k}(t)=x \right)=e^{-p\lambda\left( t \right)}\frac{{(p\lambda\left( t \right))}^{x}}{x!}=e^{\frac{ln\left( 1-{mp\mu}_{0}t \right)}{m}}\frac{{(-\frac{ln\left( 1-{mp\mu}_{0}t \right)}{m})}^{x}}{x!} Eq. (A4.2)$$

$$P\left( K(t)=x \right)=e^{-r\lambda\left( t \right)}\frac{{(r\lambda\left( t \right))}^{x}}{x!}=e^{r\frac{ln\left( 1-{mp\mu}_{0}t \right)}{mp}}\frac{{(-r\frac{ln\left( 1-{mp\mu}_{0}t \right)}{mp})}^{x}}{x!} Eq. (A4.3)$$

For the deterministic runaway model, the risk of reaching phenotypic state at age t (i.e. $K(t)\geq K_{m}$, $K_{m}$ is the threshold to reach phenotypic state) is

$$R_{0}\left( t \right)=P\left( K(t)\geq K_{m} \right)=1-\sum_{x=0}^{K_{m}-1} e^{r\lambda\left( t \right)}\frac{\left( -r\lambda\left( t \right) \right)^{x}}{x!} Eq. (A5.1)$$

$K_{m}$=5 is the cancer risk in somatic tissues; $K_{m}$=2 is the synthetic lethal risk in the germline; $K_{m}$=1 is the lethality risk in the germline.

For the probabilistic runaway model in the soma, the cancer risk at t is given by

$$R_{0}\left( t \right)=P\left( K(t)\geq10 \right)+\sum_{x=5}^{9} [P\left( K\left( t \right)=x \right)\frac{x}{12}\prod_{i=5}^{x-1} \left( 1-\frac{i}{12} \right)]=1-\sum_{x=0}^{9} e^{r\lambda\left( t \right)}\frac{\left( -r\lambda\left( t \right) \right)^{x}}{x!}+\sum_{x=5}^{9} \left[ e^{r\lambda\left( t \right)}\frac{\left( -r\lambda\left( t \right) \right)^{x}}{x!}\frac{x}{12}\prod_{i=5}^{x-1} \left( 1-\frac{i}{12} \right) \right] Eq.(A5.2)$$

We find that the analytical solutions are very close to the simulation results when the mutator effect is small (Fig. S3). The disparity grows as the mutator effect increases (Fig. S4). This is mainly because our estimate of the mutator effect mean is approximate in Eq. (A1).

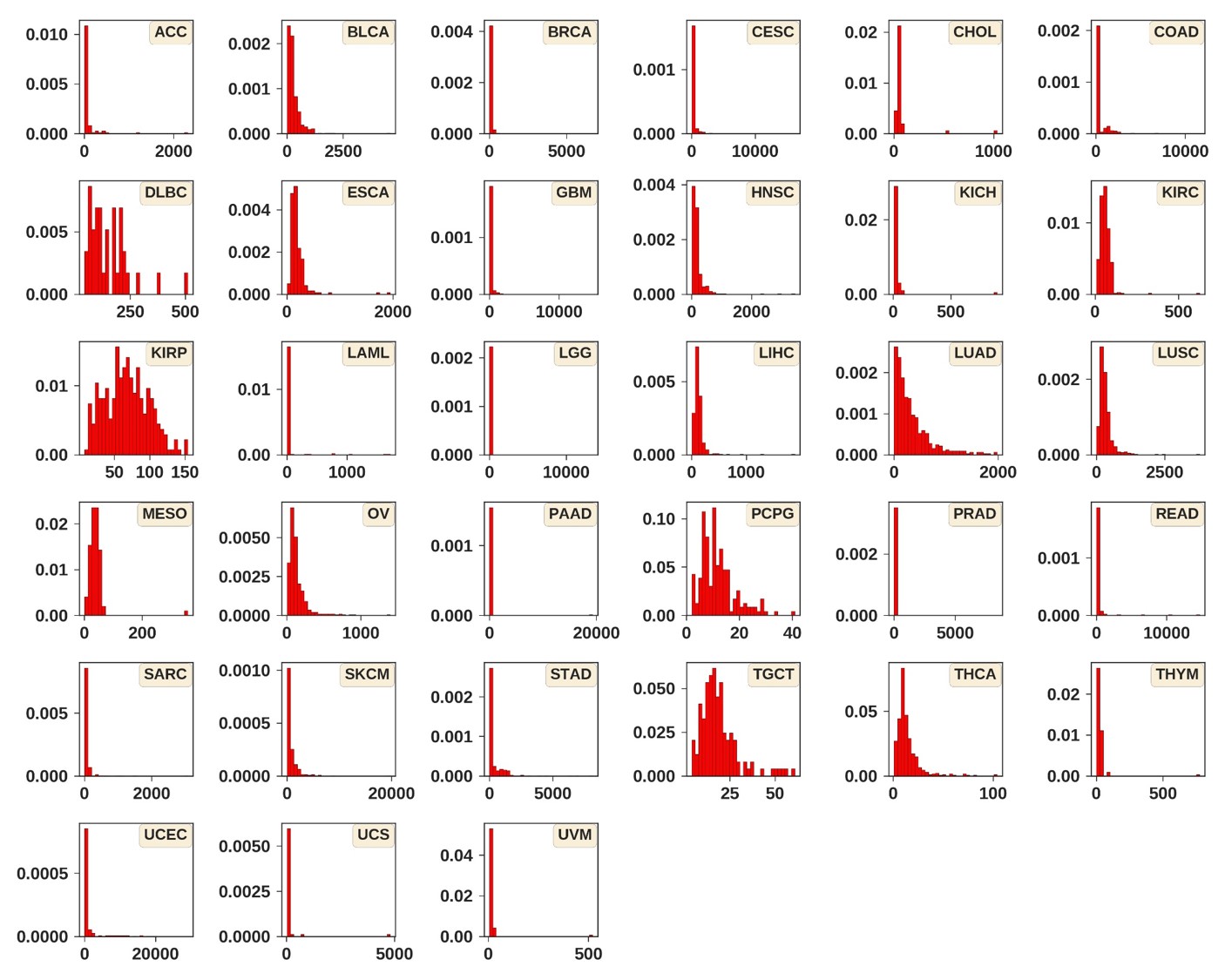

**Fig. S1. Mutation load distribution across 33 cancer types from TCGA data.** X-axis is the number of SNVs (single nucleotide variations) on exome. Y-axis is the density. There are 9979 samples across 33 cancer types.

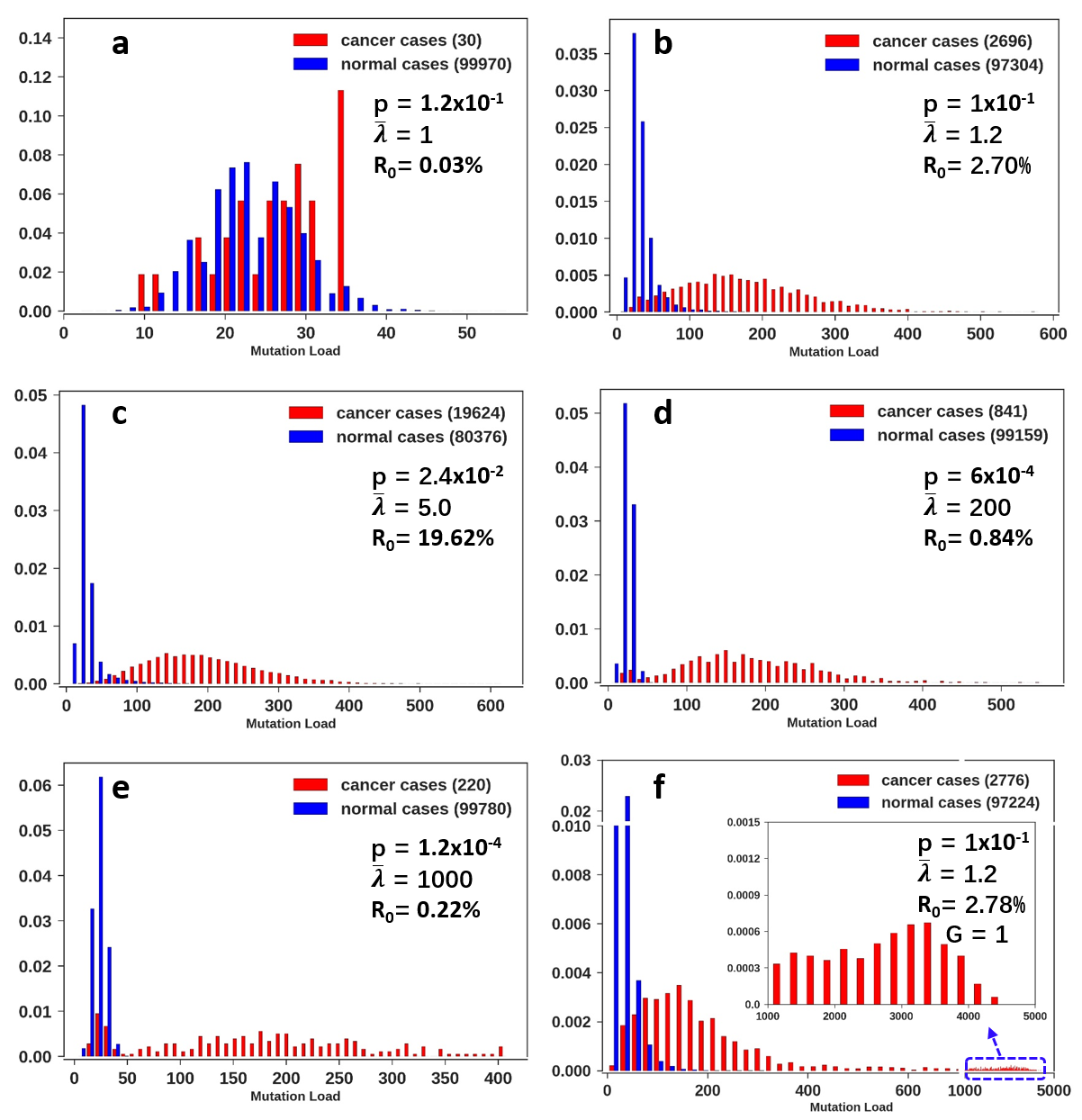

**Fig. S2. Distribution of mutational load under the probabilistic runaway model across mutator effect sizes.** (a-e) Normalized histogram of mutation load with $p\times\bar{\lambda}$ = 0.12 where p is the probability of mutator mutations and $\bar{\lambda}$ is the mean mutator strength. a – e: the mutator strength continues to increase but the mutators become less prevalent. R_0_ represents the lifetime cancer risk, the onset of cancer is based on the potential function of the probabilistic runaway model (see the main text). For each parameter set, we simulate 100000 cases. (f) The same as panel (b) except for an additional parameter G=1 (year), which is the latency time between the acquisition of the five driver mutations and the exponential expansion of the cancer cell population. During this time gap, mutation accumulation may also accelerate, thus giving rise to the strong right-skewed distribution.

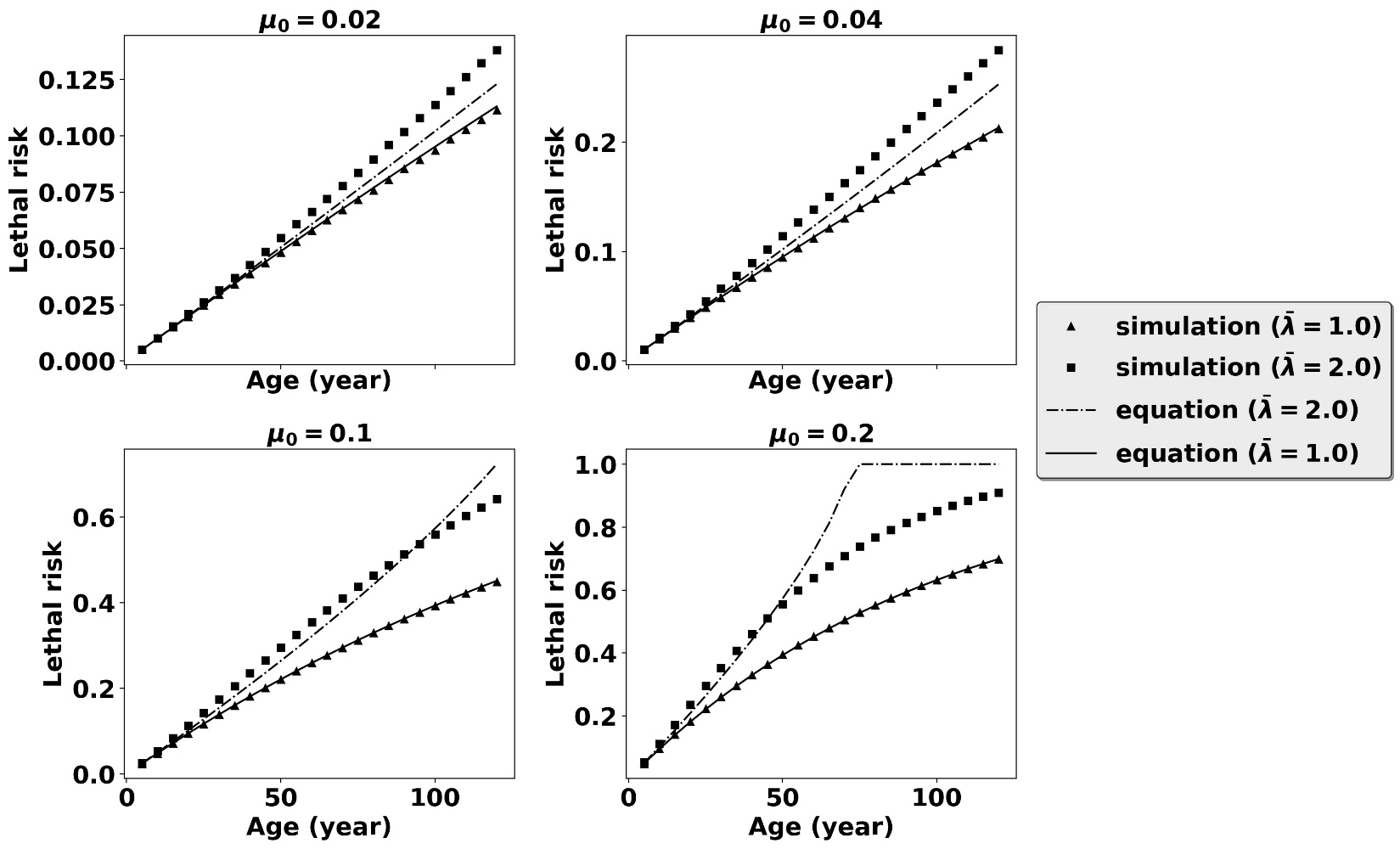

**Fig. S3. Difference between simulations and the analytical results under small mutator effect (**$\bar{\boldsymbol{\lambda}}$**=2).** The analytical results (dashed and solid lines) were obtained from Eq. (A5.1) with different initial mutation rates ($\mu_{0}$=0.02, 0.04, 0.1, and 0.2). The other common parameters are as follows if no otherwise specified: r = 0.05, p=0.1, $K_{m}$=1. Y-axis is the lethality risk of an individual suffering one or more fitness-altering mutations.

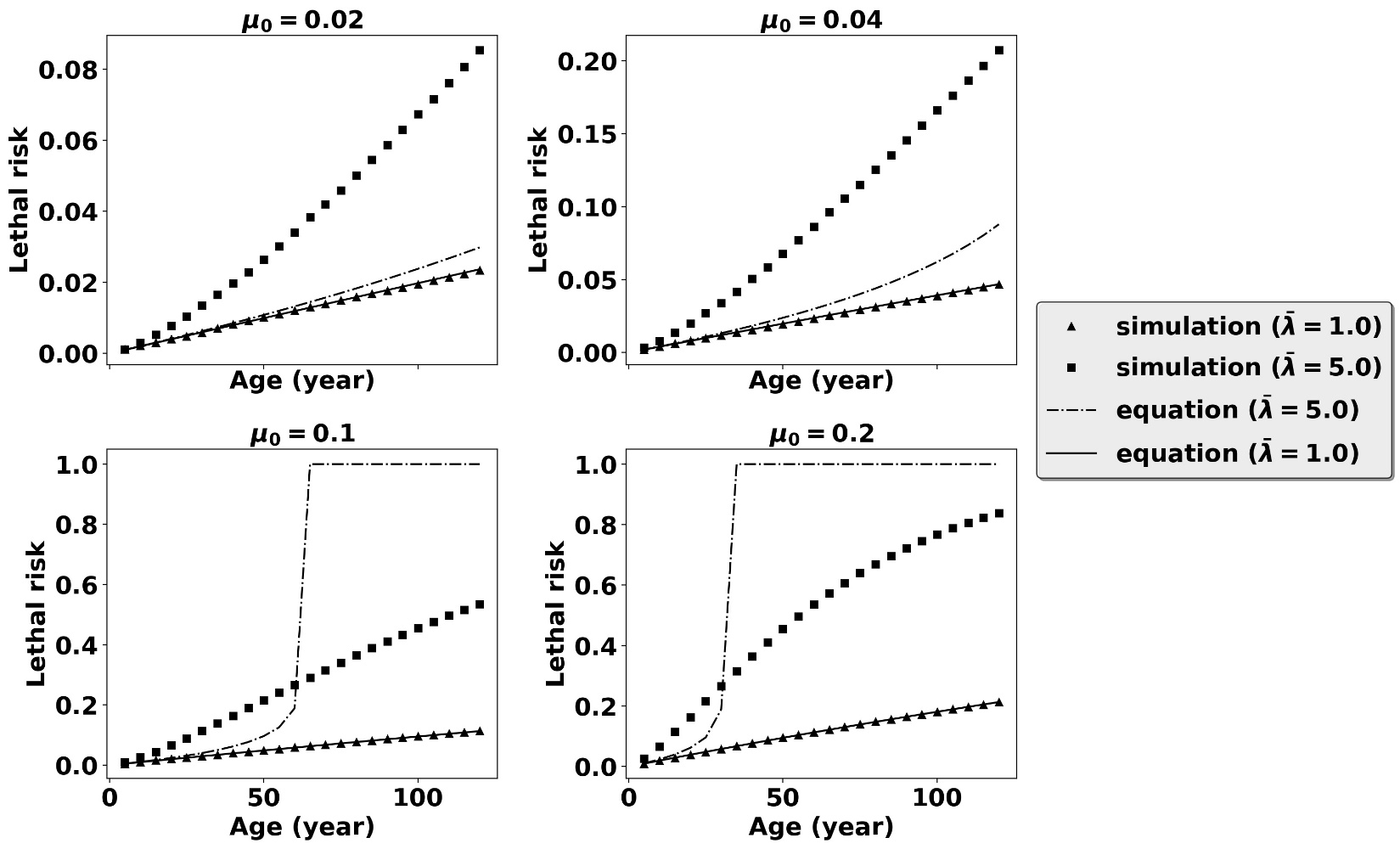

**Fig. S4. Difference between simulations and analytical results with strong mutator effect (**$\bar{\lambda}$**=5).** The analytical results (dashed and solid lines) were obtained from Eq. (A5.1) with different initial mutation rates ($\mu_{0}$=0.02, 0.04, 0.1, and 0.2). The other common parameters are as follows if no otherwise specified: r = 0.01, p=0.1, $K_{m}$=1. Y-axis is the lethality risk of an individual suffering one or more fitness-altering mutations.

**Table S1. Scheme A fitness risk (**$\bar{\boldsymbol{\lambda}}$**=2, r=0.05, p=0.1)**

| $\mu_{0}$ | 0.02 | 0.02 | 0.04 | 0.04 | 0.1 | 0.1 | 0.2 | 0.2 |
| --- | --- | --- | --- | --- | --- | --- | --- | --- |
| $\bar{\lambda}$ | 1 | 2 | 1 | 2 | 1 | 2 | 1 | 2 |
| Age |  |  |  |  |  |  |  |  |
| 5 | 0 | 0.00002 | 0.0001 | 0.00006 | 0.00036 | 0.0005 | 0.00125 | 0.0033 |
| 10 | 0.00004 | 0.0001 | 0.00019 | 0.00037 | 0.00117 | 0.00277 | 0.00471 | 0.01405 |
| 15 | 0.00009 | 0.00025 | 0.00042 | 0.00094 | 0.00251 | 0.00701 | 0.00985 | 0.03281 |
| 20 | 0.00021 | 0.00042 | 0.00076 | 0.00195 | 0.00465 | 0.01362 | 0.01696 | 0.05777 |
| 25 | 0.00031 | 0.00065 | 0.00116 | 0.00325 | 0.00743 | 0.02155 | 0.02568 | 0.08957 |
| 30 | 0.00045 | 0.00095 | 0.00178 | 0.00478 | 0.01027 | 0.03203 | 0.03607 | 0.12521 |
| 35 | 0.00059 | 0.00134 | 0.00238 | 0.00637 | 0.01374 | 0.04391 | 0.04789 | 0.16488 |
| 40 | 0.00074 | 0.00179 | 0.00306 | 0.00855 | 0.01771 | 0.05742 | 0.06096 | 0.20752 |

**Table S2. Scheme B fitness risk (**$\bar{\boldsymbol{\lambda}}$**=5, r=0.01, p=0.1)**

| $\mu_{0}$ | 0.02 | 0.02 | 0.04 | 0.04 | 0.1 | 0.1 | 0.2 | 0.2 |
| --- | --- | --- | --- | --- | --- | --- | --- | --- |
| $\bar{\lambda}$ | 1 | 5 | 1 | 5 | 1 | 5 | 1 | 5 |
| Age |  |  |  |  |  |  |  |  |
| 5 | 0 | 0.0003 | 0 | 0.00102 | 0.00002 | 0.00533 | 0.00008 | 0.01606 |
| 10 | 0 | 0.00117 | 0.00001 | 0.00334 | 0.00004 | 0.01587 | 0.00021 | 0.0455 |
| 15 | 0 | 0.00225 | 0.00001 | 0.00686 | 0.00012 | 0.02941 | 0.00043 | 0.08323 |
| 20 | 0.00001 | 0.0037 | 0.00003 | 0.01093 | 0.00025 | 0.04504 | 0.00074 | 0.12385 |
| 25 | 0.00002 | 0.00534 | 0.00004 | 0.01561 | 0.00036 | 0.06277 | 0.00114 | 0.16737 |
| 30 | 0.00002 | 0.00716 | 0.00005 | 0.0207 | 0.00051 | 0.08196 | 0.00159 | 0.21405 |
| 35 | 0.00002 | 0.00903 | 0.00006 | 0.02631 | 0.00064 | 0.10237 | 0.00217 | 0.25899 |
| 40 | 0.00004 | 0.01138 | 0.00009 | 0.03247 | 0.00083 | 0.12397 | 0.00275 | 0.30386 |

**Table S3. Exon length of 177 DNA repair genes from (Wood, et al. 2001; Wood, et al. 2005)**

| HGNC symbol | Ensembl ID | mean length | median length | longest_isoform | total length | Location |
| --- | --- | --- | --- | --- | --- | --- |
| ALKBH2 | ENSG00000189046 | 918 | 1030 | 1159 | 2290 | 12q24.11 |
| ALKBH3 | ENSG00000166199 | 999 | 764 | 1557 | 2372 | 11p11.2 |
| APEX1 | ENSG00000100823 | 929 | 831 | 1803 | 1886 | 14q11.2 |
| APEX2 | ENSG00000169188 | 2100 | 3245 | 3245 | 3245 | Xp11.21 |
| APLF | ENSG00000169621 | 2862 | 2863 | 3906 | 6553 | 2p13.3 |
| APTX | ENSG00000137074 | 1455 | 1740 | 2168 | 3789 | 9p21.1 |
| ATM | ENSG00000149311 | 7450 | 5912 | 13147 | 22317 | 11q22.3 |
| ATR | ENSG00000175054 | 5250 | 8249 | 8249 | 12496 | 3q23 |
| ATRIP | ENSG00000164053 | 2455 | 2509 | 3943 | 5111 | 3p21.31 |
| BLM | ENSG00000197299 | 3398 | 3966 | 5258 | 6551 | 15q26.1 |
| BRCA1 | ENSG00000012048 | 3151 | 2379 | 7094 | 8802 | 17q21.31 |
| BRCA2 | ENSG00000139618 | 8457 | 10984 | 11986 | 12273 | 13q13.1 |
| BRIP1 | ENSG00000136492 | 3637 | 3969 | 6048 | 7991 | 17q23.2 |
| CCNH | ENSG00000134480 | 1206 | 1378 | 2391 | 4660 | 5q14.3 |
| CDK7 | ENSG00000134058 | 915 | 818 | 1455 | 2370 | 5q13.2 |
| CETN2 | ENSG00000147400 | 1268 | 1454 | 1454 | 1928 | Xq28 |
| CHAF1A | ENSG00000167670 | 1833 | 884 | 3339 | 4299 | 19p13.3 |
| CHEK1 | ENSG00000149554 | 1957 | 1712 | 4153 | 6558 | 11q24.2 |
| CHEK2 | ENSG00000183765 | 1443 | 1545 | 2560 | 4142 | 22q12.1 |
| CLK2 | ENSG00000176444 | 1935 | 2027 | 2702 | 3670 | 1q22 |
| DCLRE1A | ENSG00000198924 | 4001 | 4241 | 4468 | 4600 | 10q25.3 |
| DCLRE1B | ENSG00000118655 | 3576 | 3576 | 3940 | 3940 | 1p13.2 |
| DCLRE1C | ENSG00000152457 | 2185 | 2463 | 3532 | 6326 | 10p13 |
| DDB1 | ENSG00000167986 | 2147 | 1375 | 4506 | 13950 | 11q12.2 |
| DDB2 | ENSG00000134574 | 1345 | 1092 | 3101 | 4897 | 11p11.2 |
| DMC1 | ENSG00000100206 | 1519 | 1595 | 2371 | 3524 | 22q13.1 |
| DUT | ENSG00000128951 | 1020 | 724 | 2147 | 3674 | 15q21.1 |
| EME1 | ENSG00000154920 | 1559 | 1538 | 2354 | 3294 | 17q21.33 |
| EME2 | ENSG00000197774 | 2630 | 1234 | 5613 | 6909 | 16p13.3 |
| ENDOV | ENSG00000173818 | 1264 | 791 | 2815 | 7253 | 17q25.3 |
| ERCC1 | ENSG00000012061 | 1319 | 938 | 3828 | 5576 | 19q13.32 |
| ERCC2 | ENSG00000104884 | 2121 | 2334 | 4153 | 6450 | 19q13.32 |
| ERCC3 | ENSG00000163161 | 2326 | 2750 | 4018 | 7401 | 2q14.3 |
| ERCC4 | ENSG00000175595 | 2805 | 2073 | 6758 | 8460 | 16p13.12 |
| ERCC5 | ENSG00000134899 | 3024 | 3660 | 5082 | 6670 | 13q33.1 |
| ERCC6 | ENSG00000225830 | 5636 | 5761 | 11026 | 14802 | 10q11.23 |
| ERCC8 | ENSG00000049167 | 1657 | 2017 | 3424 | 4847 | 5q12.1 |
| EXO1 | ENSG00000174371 | 2003 | 2899 | 3473 | 4140 | 1q43 |
| FAAP20 | ENSG00000162585 | 1888 | 2216 | 4478 | 10916 | 1p36.33 |
| FAAP24 | ENSG00000131944 | 1142 | 865 | 2298 | 2370 | 19q13.11 |
| FAN1 | ENSG00000198690 | 3490 | 4891 | 4999 | 6194 | 15q13.3 |
| FANCA | ENSG00000187741 | 2220 | 1088 | 5451 | 9736 | 16q24.3 |
| FANCB | ENSG00000181544 | 2721 | 2894 | 3008 | 3215 | Xp22.2 |
| FANCC | ENSG00000158169 | 2491 | 2709 | 4585 | 7912 | 9q22.32 |
| FANCD2 | ENSG00000144554 | 4073 | 5102 | 5219 | 8378 | 3p25.3 |
| FANCE | ENSG00000112039 | 2554 | 2554 | 2554 | 2554 | 6p21.31 |
| FANCF | ENSG00000183161 | 4269 | 4269 | 4269 | 4269 | 11p14.3 |
| FANCG | ENSG00000221829 | 1711 | 2108 | 2631 | 3614 | 9p13.3 |
| FANCI | ENSG00000140525 | 3219 | 3690 | 4743 | 7766 | 15q26.1 |
| FANCL | ENSG00000115392 | 1161 | 964 | 1698 | 2412 | 2p16.1 |
| FANCM | ENSG00000187790 | 4629 | 5577 | 7111 | 8468 | 14q21.2 |
| FEN1 | ENSG00000168496 | 1280 | 776 | 2296 | 2478 | 11q12.2 |
| GEN1 | ENSG00000178295 | 4771 | 6367 | 9854 | 11827 | 2p24.2 |
| GTF2H1 | ENSG00000110768 | 1843 | 1822 | 2982 | 6939 | 11p15.1 |
| GTF2H2 | ENSG00000145736 | 1547 | 1874 | 2077 | 6389 | 5q13.2 |
| GTF2H3 | ENSG00000111358 | 1272 | 790 | 3329 | 3996 | 12q24.31 |
| GTF2H4 | ENSG00000213780 | 1366 | 1673 | 1735 | 2353 | 6p21.33 |
| GTF2H5 | ENSG00000272047 | 7481 | 7481 | 7481 | 7481 | 6q25.3 |
| H2AFX | ENSG00000188486 | 1453 | 1373 | 1614 | 1614 | 11q23.3 |
| HELQ | ENSG00000163312 | 3070 | 3429 | 3579 | 3713 | 4q21.23 |
| HLTF | ENSG00000071794 | 3792 | 3896 | 5317 | 6024 | 3q24 |
| HUS1 | ENSG00000136273 | 1392 | 1125 | 2935 | 4177 | 7p12.3 |
| LIG1 | ENSG00000105486 | 2734 | 3179 | 4102 | 6576 | 19q13.33 |
| LIG3 | ENSG00000005156 | 2911 | 1699 | 8400 | 12919 | 17q12 |
| LIG4 | ENSG00000174405 | 4035 | 4065 | 4180 | 4387 | 13q33.3 |
| MAD2L2 | ENSG00000116670 | 1086 | 1002 | 1860 | 3324 | 1p36.22 |
| MBD4 | ENSG00000129071 | 1714 | 2099 | 2478 | 3356 | 3q21.3 |
| MDC1 | ENSG00000137337 | 3007 | 916 | 7576 | 8064 | 6p21.33 |
| MGMT | ENSG00000170430 | 1113 | 867 | 1759 | 3557 | 10q26.3 |
| MLH1 | ENSG00000076242 | 1680 | 2230 | 2752 | 3532 | 3p22.2 |
| MLH3 | ENSG00000119684 | 3613 | 2118 | 7896 | 8363 | 14q24.3 |
| MMS19 | ENSG00000155229 | 2480 | 3165 | 3797 | 5849 | 10q24.1 |
| MNAT1 | ENSG00000020426 | 1131 | 680 | 2635 | 3865 | 14q23.1 |
| MPG | ENSG00000103152 | 1026 | 1039 | 1205 | 1661 | 16p13.3 |
| MPLKIP | ENSG00000168303 | 7668 | 7668 | 7668 | 7668 | 7p14.1 |
| MRE11 | ENSG00000020922 | 2887 | 2604 | 6897 | 7689 | 11q21 |
| MSH2 | ENSG00000095002 | 2898 | 2918 | 3628 | 4552 | 2p21-p16.3 |
| MSH3 | ENSG00000113318 | 3229 | 4092 | 4092 | 4709 | 5q14.1 |
| MSH4 | ENSG00000057468 | 3266 | 3266 | 3266 | 3266 | 1p31.1 |
| MSH5 | ENSG00000204410 | 2110 | 2669 | 3094 | 6234 | 6p21.33 |
| MSH6 | ENSG00000116062 | 3506 | 4055 | 7476 | 10993 | 2p16.3 |
| MUS81 | ENSG00000172732 | 1545 | 1793 | 3420 | 5476 | 11q13 |
| MUTYH | ENSG00000132781 | 1366 | 1677 | 1929 | 3909 | 1p34.1 |
| NABP2 | ENSG00000139579 | 1173 | 1411 | 1775 | 2218 | 12q13.3 |
| NBN | ENSG00000104320 | 3101 | 4523 | 4907 | 6156 | 8q21.3 |
| NEIL1 | ENSG00000140398 | 1461 | 833 | 3759 | 6938 | 15q24.2 |
| NEIL2 | ENSG00000154328 | 1827 | 2226 | 2671 | 3367 | 8p23.1 |
| NEIL3 | ENSG00000109674 | 2156 | 2408 | 2408 | 2959 | 4q34.3 |
| NHEJ1 | ENSG00000187736 | 1243 | 753 | 2113 | 3109 | 2q35 |
| NTHL1 | ENSG00000065057 | 803 | 814 | 1067 | 2430 | 16p13.3 |
| NUDT1 | ENSG00000106268 | 716 | 714 | 872 | 2093 | 7p22.3 |
| OGG1 | ENSG00000114026 | 1517 | 1699 | 4977 | 8733 | 3p25.3 |
| PALB2 | ENSG00000083093 | 2714 | 3963 | 4003 | 4747 | 16p12.2 |
| PARP1 | ENSG00000143799 | 2408 | 3165 | 3958 | 7165 | 1q42.12 |
| PARP2 | ENSG00000129484 | 1420 | 1881 | 1980 | 4096 | 14q11.2 |
| PARP3 | ENSG00000041880 | 1877 | 2336 | 2472 | 2928 | 3p21.2 |
| PCNA | ENSG00000132646 | 1352 | 1359 | 1359 | 1471 | 20p12.3 |
| PER1 | ENSG00000179094 | 2908 | 3209 | 4707 | 7002 | 17p13.1 |
| PMS1 | ENSG00000064933 | 2363 | 2795 | 3417 | 5448 | 2q32.2 |
| PMS2 | ENSG00000122512 | 2009 | 1719 | 2855 | 3936 | 7p22.1 |
| PNKP | ENSG00000039650 | 1436 | 1633 | 2102 | 3531 | 19q13.33 |
| POLB | ENSG00000070501 | 884 | 744 | 1341 | 4106 | 8p11.21 |
| POLD1 | ENSG00000062822 | 2711 | 3435 | 3542 | 4858 | 19q13.3 |
| POLE | ENSG00000177084 | 5540 | 6800 | 8011 | 16609 | 12q24.33 |
| POLG | ENSG00000140521 | 2670 | 4487 | 4502 | 5529 | 15q26.1 |
| POLH | ENSG00000170734 | 3059 | 3322 | 3540 | 3540 | 6p21.1 |
| POLI | ENSG00000101751 | 2011 | 1734 | 6133 | 9471 | 18q21.2 |
| POLK | ENSG00000122008 | 2669 | 2482 | 5911 | 9105 | 5q13.3 |
| POLL | ENSG00000166169 | 1683 | 1457 | 2914 | 5420 | 10q24.32 |
| POLM | ENSG00000122678 | 1813 | 2380 | 3218 | 4487 | 7p13 |
| POLN | ENSG00000130997 | 2335 | 2899 | 3978 | 7721 | 4p16.3 |
| POLQ | ENSG00000051341 | 7760 | 8775 | 9055 | 9400 | 3q13.33 |
| PRKDC | ENSG00000253729 | 11291 | 12784 | 13509 | 15417 | 8q11.21 |
| PRPF19 | ENSG00000110107 | 1206 | 943 | 2157 | 2739 | 11q12.2 |
| RAD1 | ENSG00000113456 | 2145 | 1328 | 5770 | 6763 | 5p13.2 |
| RAD17 | ENSG00000152942 | 2372 | 2789 | 3233 | 4159 | 5q13.2 |
| RAD18 | ENSG00000070950 | 1917 | 592 | 5886 | 6983 | 3p25.3 |
| RAD23A | ENSG00000179262 | 1359 | 1526 | 1746 | 2449 | 19p13.13 |
| RAD23B | ENSG00000119318 | 2672 | 3835 | 4119 | 4615 | 9q31.2 |
| RAD50 | ENSG00000113522 | 4057 | 2915 | 8306 | 9771 | 5q31.1 |
| RAD51 | ENSG00000051180 | 1400 | 1530 | 2449 | 3046 | 15q15.1 |
| RAD51B | ENSG00000182185 | 1115 | 829 | 2596 | 7477 | 14q24.1 |
| RAD51C | ENSG00000108384 | 1154 | 779 | 2591 | 6068 | 17q22 |
| RAD51D | ENSG00000185379 | 1748 | 1194 | 9988 | 11304 | 17q12 |
| RAD52 | ENSG00000002016 | 1670 | 1530 | 2826 | 4190 | 12p13.33 |
| RAD54B | ENSG00000197275 | 2107 | 1879 | 3089 | 5763 | 8q22.1 |
| RAD54L | ENSG00000085999 | 1513 | 822 | 3108 | 3212 | 1p34.1 |
| RAD9A | ENSG00000172613 | 1117 | 924 | 2119 | 4169 | 11q13.2 |
| RBBP8 | ENSG00000101773 | 2119 | 2962 | 3293 | 5091 | 18q11.2 |
| RDM1 | ENSG00000278023 | 832 | 686 | 1535 | 1975 | 17q12 |
| RECQL | ENSG00000004700 | 1921 | 1054 | 3702 | 3778 | 12p12.1 |
| RECQL4 | ENSG00000160957 | 2368 | 2487 | 3896 | 4236 | 8q24.3 |
| RECQL5 | ENSG00000108469 | 2288 | 1972 | 3710 | 8766 | 17q25 |
| REV1 | ENSG00000135945 | 3254 | 4686 | 4751 | 7769 | 2q11.2 |
| REV3L | ENSG00000009413 | 9421 | 10701 | 10964 | 12425 | 6q21 |
| RIF1 | ENSG00000080345 | 7706 | 7897 | 15003 | 18166 | 2q23.3 |
| RNF168 | ENSG00000163961 | 3917 | 5347 | 5347 | 5347 | 3q29 |
| RNF4 | ENSG00000063978 | 1294 | 615 | 4123 | 7270 | 4p16.3 |
| RNF8 | ENSG00000112130 | 2205 | 1800 | 5631 | 6575 | 6p21.2 |
| RPA1 | ENSG00000132383 | 2354 | 844 | 4340 | 5492 | 17p13.3 |
| RPA2 | ENSG00000117748 | 1161 | 1162 | 1752 | 2019 | 1p35.3 |
| RPA3 | ENSG00000106399 | 1060 | 847 | 2020 | 3232 | 7p21.3 |
| RPA4 | ENSG00000204086 | 1560 | 1560 | 1560 | 1560 | Xq21.33 |
| RRM2B | ENSG00000048392 | 1908 | 900 | 4931 | 5548 | 8q22.3 |
| SEM1 | ENSG00000127922 | 1008 | 610 | 2773 | 11490 | 7q21.3 |
| SETMAR | ENSG00000170364 | 1410 | 1657 | 2443 | 3567 | 3p26.1 |
| SHPRH | ENSG00000146414 | 6460 | 6649 | 11660 | 17637 | 6q24.3 |
| SLX1A | ENSG00000132207 | 997 | 1091 | 1558 | 2464 | 16p11.2 |
| SLX1B | ENSG00000181625 | 1050 | 1165 | 1558 | 2462 | 16p11.2 |
| SLX4 | ENSG00000188827 | 5557 | 7307 | 7307 | 8591 | 16p13.3 |
| SMUG1 | ENSG00000123415 | 941 | 627 | 2607 | 4981 | 12q13.13 |
| SPO11 | ENSG00000054796 | 1465 | 1608 | 1834 | 1867 | 20q13.31 |
| SPRTN | ENSG00000010072 | 1936 | 1667 | 3559 | 5906 | 1q42.2 |
| TDG | ENSG00000139372 | 1511 | 1277 | 3251 | 4903 | 12q23.3 |
| TDP1 | ENSG00000042088 | 2076 | 2200 | 3702 | 6227 | 14q32.11 |
| TDP2 | ENSG00000111802 | 1523 | 2071 | 2120 | 3082 | 6p22.3 |
| TOPBP1 | ENSG00000163781 | 2899 | 1142 | 5378 | 6515 | 3q22.1 |
| TP53 | ENSG00000141510 | 2045 | 2506 | 2724 | 3936 | 17p13.1 |
| TP53BP1 | ENSG00000067369 | 5318 | 6015 | 10384 | 12991 | 15q15.3 |
| TREX1 | ENSG00000213689 | 1299 | 1105 | 2197 | 2197 | 3p21.31 |
| TREX2 | ENSG00000183479 | 1979 | 2028 | 2262 | 2715 | Xq28 |
| UBE2A | ENSG00000077721 | 1352 | 1796 | 2643 | 3251 | Xq24 |
| UBE2B | ENSG00000119048 | 921 | 644 | 2515 | 3291 | 5q31.1 |
| UBE2N | ENSG00000177889 | 1854 | 960 | 5186 | 6269 | 12q22 |
| UBE2V2 | ENSG00000169139 | 1496 | 918 | 4373 | 6606 | 8q11.21 |
| UNG | ENSG00000076248 | 1946 | 2071 | 2234 | 2567 | 12q24.11 |
| UVSSA | ENSG00000163945 | 2920 | 2927 | 4819 | 7830 | 4p16.3 |
| WRN | ENSG00000165392 | 4377 | 5215 | 5215 | 5634 | 8p12 |
| XAB2 | ENSG00000076924 | 2974 | 2667 | 4555 | 4673 | 19p13.3 |
| XPA | ENSG00000136936 | 1195 | 1417 | 1584 | 1833 | 9q22.33 |
| XPC | ENSG00000154767 | 2275 | 2944 | 3832 | 4798 | 3p25.1 |
| XRCC1 | ENSG00000073050 | 1506 | 1147 | 2802 | 3459 | 19q13.2 |
| XRCC2 | ENSG00000196584 | 3897 | 3897 | 4728 | 4853 | 7q36.1 |
| XRCC3 | ENSG00000126215 | 1690 | 2539 | 3668 | 7507 | 14q32.33 |
| XRCC4 | ENSG00000152422 | 1540 | 1579 | 1696 | 2676 | 5q14.2 |
| XRCC5 | ENSG00000079246 | 2531 | 2962 | 3761 | 5463 | 2q35 |
| XRCC6 | ENSG00000196419 | 2088 | 2148 | 2709 | 3288 | 22q13.2 |

**Table S4. Exon length of 299 cancer driver genes from (Lawrence, et al. 2014; Bailey, et al. 2018)**

| HGNC symbol | Ensembl ID | mean length | median length | longest_isoform | total length | Location |
| --- | --- | --- | --- | --- | --- | --- |
| ABL1 | ENSG00000097007 | 4439 | 3824 | 5766 | 6276 | 9q34.12 |
| ACVR1 | ENSG00000115170 | 1899 | 2084 | 3047 | 3990 | 2q24.1 |
| ACVR1B | ENSG00000135503 | 2191 | 1791 | 4563 | 5690 | 12q13.13 |
| ACVR2A | ENSG00000121989 | 3473 | 2367 | 5728 | 6770 | 2q22.3-q23.1 |
| AJUBA | ENSG00000129474 | 1900 | 2569 | 4262 | 5979 | 14q11.2 |
| AKT1 | ENSG00000142208 | 2422 | 2710 | 8396 | 11162 | 14q32.33 |
| ALB | ENSG00000163631 | 1431 | 1519 | 2263 | 3377 | 4q13.3 |
| ALK | ENSG00000171094 | 4411 | 4646 | 6220 | 6932 | 2p23.2-p23.1 |
| AMER1 | ENSG00000184675 | 5590 | 3688 | 8443 | 8443 | Xq11.2 |
| APC | ENSG00000134982 | 6141 | 5733 | 10701 | 12440 | 5q22.2 |
| APOB | ENSG00000084674 | 11610 | 13991 | 14121 | 14828 | 2p24.1 |
| AR | ENSG00000169083 | 6137 | 3899 | 10676 | 12775 | Xq12 |
| ARAF | ENSG00000078061 | 1929 | 2458 | 2504 | 3535 | Xp11.3 |
| ARHGAP35 | ENSG00000160007 | 5129 | 7696 | 8889 | 11245 | 19q13.32 |
| ARID1A | ENSG00000117713 | 5357 | 5992 | 8577 | 9246 | 1p36.11 |
| ARID2 | ENSG00000189079 | 6096 | 7316 | 8642 | 12127 | 12q12 |
| ARID5B | ENSG00000150347 | 5935 | 7891 | 7891 | 8164 | 10q21.2 |
| ASXL1 | ENSG00000171456 | 4079 | 5374 | 7038 | 9454 | 20q11.21 |
| ASXL2 | ENSG00000143970 | 7745 | 8889 | 12878 | 13023 | 2p23.3 |
| ATF7IP | ENSG00000171681 | 4005 | 4142 | 8878 | 13189 | 12p13.1 |
| ATM | ENSG00000149311 | 7450 | 5912 | 13147 | 22317 | 11q22.3 |
| ATR | ENSG00000175054 | 5250 | 8249 | 8249 | 12496 | 3q23 |
| ATRX | ENSG00000085224 | 6625 | 10218 | 11220 | 19060 | Xq21.1 |
| ATXN3 | ENSG00000066427 | 2011 | 924 | 26812 | 28202 | 14q32.12 |
| AXIN1 | ENSG00000103126 | 3526 | 3643 | 6340 | 7487 | 16p13.3 |
| AXIN2 | ENSG00000168646 | 2784 | 2541 | 4259 | 5226 | 17q24.1 |
| B2M | ENSG00000166710 | 841 | 717 | 2329 | 3728 | 15q21.1 |
| BAP1 | ENSG00000163930 | 2159 | 1654 | 3937 | 5055 | 3p21.1 |
| BCL2 | ENSG00000171791 | 3325 | 3209 | 7461 | 7665 | 18q21.33 |
| BCL2L11 | ENSG00000153094 | 2434 | 947 | 5231 | 6658 | 2q13 |
| BCOR | ENSG00000183337 | 4603 | 6258 | 6390 | 8544 | Xp11.4 |
| BRAF | ENSG00000157764 | 3207 | 2408 | 8294 | 10380 | 7q34 |
| BRCA1 | ENSG00000012048 | 3151 | 2379 | 7094 | 8802 | 17q21.31 |
| BRCA2 | ENSG00000139618 | 8457 | 10984 | 11986 | 12273 | 13q13.1 |
| BRD7 | ENSG00000166164 | 2636 | 2145 | 5370 | 8492 | 16q12.1 |
| BTG2 | ENSG00000159388 | 1849 | 1275 | 2712 | 2757 | 1q32.1 |
| CACNA1A | ENSG00000141837 | 6106 | 7810 | 8410 | 10920 | 19p13.13 |
| CARD11 | ENSG00000198286 | 3013 | 4366 | 4366 | 4707 | 7p22.2 |
| CASP8 | ENSG00000064012 | 1536 | 1629 | 2930 | 7243 | 2q33.1 |
| CBFB | ENSG00000067955 | 1482 | 728 | 3136 | 4129 | 16q22.1 |
| CBWD3 | ENSG00000196873 | 1877 | 1813 | 3301 | 12319 | 9q21.11 |
| CCND1 | ENSG00000110092 | 1771 | 744 | 4307 | 4830 | 11q13.3 |
| CD70 | ENSG00000125726 | 777 | 881 | 913 | 1532 | 19p13.3 |
| CD79B | ENSG00000007312 | 967 | 1206 | 1269 | 2080 | 17q23.3 |
| CDH1 | ENSG00000039068 | 3048 | 2759 | 4889 | 6177 | 16q22.1 |
| CDK12 | ENSG00000167258 | 4350 | 4165 | 8336 | 12773 | 17q12 |
| CDK4 | ENSG00000135446 | 915 | 825 | 2076 | 3168 | 12q14.1 |
| CDKN1A | ENSG00000124762 | 1474 | 2102 | 2267 | 3050 | 6p21.2 |
| CDKN1B | ENSG00000111276 | 1109 | 577 | 2657 | 2811 | 12p13.1 |
| CDKN2A | ENSG00000147889 | 853 | 856 | 1283 | 4329 | 9p21.3 |
| CDKN2C | ENSG00000123080 | 2257 | 2086 | 2904 | 3318 | 1p32.3 |
| CEBPA | ENSG00000245848 | 2631 | 2631 | 2631 | 2631 | 19q13.11 |
| CHD3 | ENSG00000170004 | 5208 | 7286 | 7356 | 9758 | 17p13.1 |
| CHD4 | ENSG00000111642 | 5036 | 6496 | 6554 | 7474 | 12p13.31 |
| CHD8 | ENSG00000100888 | 5833 | 7420 | 8229 | 9781 | 14q11.2 |
| CHEK2 | ENSG00000183765 | 1443 | 1545 | 2560 | 4142 | 22q12.1 |
| CIC | ENSG00000079432 | 5151 | 5792 | 8218 | 9494 | 19q13.2 |
| CNBD1 | ENSG00000176571 | 907 | 659 | 1594 | 2584 | 8q21.3 |
| CNOT9 | ENSG00000144580 | 1958 | 1129 | 4079 | 5196 | 2q35 |
| COL5A1 | ENSG00000130635 | 7346 | 8437 | 8471 | 11189 | 9q34.3 |
| CREB3L3 | ENSG00000060566 | 1592 | 1475 | 2618 | 2678 | 19p13.3 |
| CREBBP | ENSG00000005339 | 5893 | 7598 | 10803 | 15068 | 16p13.3 |
| CSDE1 | ENSG00000009307 | 3167 | 3774 | 4312 | 5778 | 1p13.2 |
| CTCF | ENSG00000102974 | 3277 | 2978 | 3939 | 4301 | 16q22.1 |
| CTNNB1 | ENSG00000168036 | 2410 | 2841 | 3737 | 6614 | 3p22.1 |
| CTNND1 | ENSG00000198561 | 5508 | 5832 | 6553 | 8833 | 11q12.1 |
| CUL1 | ENSG00000055130 | 2870 | 2955 | 3064 | 3548 | 7q36.1 |
| CUL3 | ENSG00000036257 | 3123 | 2701 | 6741 | 11505 | 2q36.2 |
| CYLD | ENSG00000083799 | 3862 | 3513 | 8503 | 12954 | 16q12.1 |
| CYSLTR2 | ENSG00000152207 | 1173 | 727 | 4672 | 5372 | 13q14.2 |
| DACH1 | ENSG00000276644 | 4318 | 4708 | 5233 | 5389 | 13q21.33 |
| DAZAP1 | ENSG00000071626 | 2016 | 2157 | 3943 | 9201 | 19p13.3 |
| DDX3X | ENSG00000215301 | 2657 | 1592 | 5399 | 8881 | Xp11.4 |
| DHX9 | ENSG00000135829 | 2567 | 2531 | 4240 | 5645 | 1q25.3 |
| DIAPH2 | ENSG00000147202 | 5028 | 3782 | 9333 | 9550 | Xq21.33 |
| DICER1 | ENSG00000100697 | 6678 | 6179 | 10331 | 12294 | 14q32.13 |
| DMD | ENSG00000198947 | 9150 | 7410 | 13956 | 18239 | Xp21.2-p21.1 |
| DNMT3A | ENSG00000119772 | 3662 | 3589 | 9501 | 12020 | 2p23.3 |
| EEF1A1 | ENSG00000156508 | 2377 | 2303 | 4441 | 5948 | 6q13 |
| EEF2 | ENSG00000167658 | 1749 | 581 | 3164 | 4027 | 19p13.3 |
| EGFR | ENSG00000146648 | 4421 | 3844 | 9821 | 12961 | 7p11.2 |
| EGR3 | ENSG00000179388 | 1795 | 1500 | 4336 | 4855 | 8p21.3 |
| EIF1AX | ENSG00000173674 | 2736 | 4427 | 4427 | 4427 | Xp22.12 |
| ELF3 | ENSG00000163435 | 2096 | 2044 | 4994 | 7191 | 1q32.1 |
| ELOC | ENSG00000154582 | 1294 | 816 | 2780 | 4934 | 8q21.11 |
| EP300 | ENSG00000100393 | 9585 | 9585 | 9585 | 9585 | 22q13.2 |
| EPAS1 | ENSG00000116016 | 2390 | 1067 | 5166 | 7302 | 2p21 |
| EPHA2 | ENSG00000142627 | 3230 | 3963 | 3964 | 4164 | 1p36.13 |
| EPHA3 | ENSG00000044524 | 4122 | 3085 | 5809 | 6674 | 3p11.1 |
| ERBB2 | ENSG00000141736 | 3457 | 4341 | 4930 | 10321 | 17q12 |
| ERBB3 | ENSG00000065361 | 3210 | 3289 | 5919 | 8658 | 12q13.2 |
| ERBB4 | ENSG00000178568 | 7987 | 11755 | 12136 | 13023 | 2q34 |
| ERCC2 | ENSG00000104884 | 2121 | 2334 | 4153 | 6450 | 19q13.32 |
| ESR1 | ENSG00000091831 | 3843 | 4385 | 6466 | 13629 | 6q25.1-q25.2 |
| EZH2 | ENSG00000106462 | 2527 | 2522 | 3641 | 4522 | 7q36.1 |
| FAT1 | ENSG00000083857 | 9820 | 14758 | 14786 | 16177 | 4q35.2 |
| FBXW7 | ENSG00000109670 | 2669 | 2562 | 4976 | 10287 | 4q31.3 |
| FGFR1 | ENSG00000077782 | 3074 | 2953 | 5900 | 12698 | 8p11.23 |
| FGFR2 | ENSG00000066468 | 2988 | 3003 | 4369 | 7486 | 10q26.13 |
| FGFR3 | ENSG00000068078 | 3863 | 4217 | 4438 | 4834 | 4p16.3 |
| FLNA | ENSG00000196924 | 7220 | 8225 | 8486 | 10585 | Xq28 |
| FLT3 | ENSG00000122025 | 3547 | 3634 | 3842 | 4159 | 13q12.2 |
| FOXA1 | ENSG00000129514 | 1201 | 789 | 2862 | 3982 | 14q21.1 |
| FOXA2 | ENSG00000125798 | 2409 | 2401 | 2422 | 2603 | 20p11.21 |
| FOXQ1 | ENSG00000164379 | 1715 | 1715 | 1715 | 1715 | 6p25.3 |
| FUBP1 | ENSG00000162613 | 1721 | 2201 | 2524 | 5666 | 1p31.1 |
| GABRA6 | ENSG00000145863 | 1291 | 591 | 2393 | 3738 | 5q34 |
| GATA3 | ENSG00000107485 | 2233 | 2654 | 3078 | 3253 | 10p14 |
| GNA11 | ENSG00000088256 | 1698 | 736 | 4147 | 6242 | 19p13.3 |
| GNA13 | ENSG00000120063 | 3212 | 3212 | 4922 | 5104 | 17q24.1 |
| GNAQ | ENSG00000156052 | 4361 | 6539 | 6539 | 6629 | 9q21.2 |
| GNAS | ENSG00000087460 | 1449 | 911 | 4029 | 13320 | 20q13.32 |
| GPS2 | ENSG00000132522 | 1196 | 1238 | 1977 | 3089 | 17p13.1 |
| GRIN2D | ENSG00000105464 | 5093 | 5093 | 5093 | 5093 | 19q13.33 |
| GTF2I | ENSG00000263001 | 3224 | 4352 | 4548 | 10438 | 7q11.23 |
| H3F3A | ENSG00000163041 | 1190 | 1088 | 1863 | 3146 | 1q42.12 |
| H3F3C | ENSG00000188375 | 1057 | 1057 | 1057 | 1057 | 12p11.21 |
| HGF | ENSG00000019991 | 2471 | 1990 | 5989 | 8897 | 7q21.11 |
| HIST1H1C | ENSG00000187837 | 642 | 642 | 642 | 642 | 6p22.2 |
| HIST1H1E | ENSG00000168298 | 660 | 660 | 660 | 660 | 6p22.2 |
| HLA-A | ENSG00000206503 | 1641 | 1611 | 1868 | 2710 | 6p22.1 |
| HLA-B | ENSG00000234745 | 1127 | 1087 | 1547 | 3012 | 6p21.33 |
| HRAS | ENSG00000174775 | 933 | 1099 | 1233 | 1854 | 11p15.5 |
| HUWE1 | ENSG00000086758 | 11468 | 14311 | 14796 | 17380 | Xp11.22 |
| IDH1 | ENSG00000138413 | 1693 | 2298 | 2441 | 4721 | 2q34 |
| IDH2 | ENSG00000182054 | 1766 | 1453 | 2694 | 2814 | 15q26.1 |
| IL6ST | ENSG00000134352 | 4302 | 3007 | 9057 | 9292 | 5q11.2 |
| IL7R | ENSG00000168685 | 1553 | 917 | 4626 | 6060 | 5p13.2 |
| INPPL1 | ENSG00000165458 | 3236 | 4649 | 4733 | 6519 | 11q13.4 |
| IRF2 | ENSG00000168310 | 876 | 591 | 2303 | 3116 | 4q35.1 |
| IRF6 | ENSG00000117595 | 3162 | 4256 | 4306 | 4915 | 1q32.2 |
| JAK1 | ENSG00000162434 | 3274 | 5047 | 5047 | 6317 | 1p31.3 |
| JAK2 | ENSG00000096968 | 4488 | 5285 | 5285 | 5794 | 9p24.1 |
| JAK3 | ENSG00000105639 | 3480 | 3612 | 5432 | 6813 | 19p13.11 |
| KANSL1 | ENSG00000120071 | 4145 | 5014 | 9095 | 13928 | 17q21.31 |
| KDM5C | ENSG00000126012 | 4615 | 5245 | 6096 | 8727 | Xp11.22 |
| KDM6A | ENSG00000147050 | 4695 | 5633 | 5924 | 7139 | Xp11.3 |
| KEAP1 | ENSG00000079999 | 1523 | 986 | 2955 | 3761 | 19p13.2 |
| KEL | ENSG00000197993 | 1515 | 786 | 2812 | 3923 | 7q34 |
| KIF1A | ENSG00000130294 | 5830 | 5533 | 9223 | 12357 | 2q37.3 |
| KIT | ENSG00000157404 | 3871 | 3470 | 5186 | 5563 | 4q12 |
| KLF5 | ENSG00000102554 | 2130 | 2951 | 3566 | 3893 | 13q22.1 |
| KMT2A | ENSG00000118058 | 9762 | 13655 | 16602 | 21961 | 11q23.3 |
| KMT2B | ENSG00000272333 | 6256 | 8469 | 8469 | 10279 | 19q13.12 |
| KMT2C | ENSG00000055609 | 13600 | 16858 | 16862 | 20114 | 7q36.1 |
| KMT2D | ENSG00000167548 | 14361 | 19419 | 19419 | 20476 | 12q13.12 |
| KRAS | ENSG00000133703 | 2575 | 1119 | 5765 | 7302 | 12p12.1 |
| KRT222 | ENSG00000213424 | 1725 | 1712 | 2764 | 3169 | 17q21.2 |
| LATS1 | ENSG00000131023 | 3859 | 3879 | 7517 | 8643 | 6q25.1 |
| LATS2 | ENSG00000150457 | 4143 | 5511 | 5511 | 5578 | 13q12.11 |
| LEMD2 | ENSG00000161904 | 1727 | 1306 | 4870 | 6788 | 6p21.31 |
| LZTR1 | ENSG00000099949 | 2379 | 2323 | 4572 | 7556 | 22q11.21 |
| MACF1 | ENSG00000127603 | 15753 | 17538 | 24828 | 43102 | 1p34.3 |
| MAP2K1 | ENSG00000169032 | 2357 | 1432 | 3410 | 3759 | 15q22.31 |
| MAP2K4 | ENSG00000065559 | 1610 | 977 | 3856 | 5714 | 17p12 |
| MAP3K1 | ENSG00000095015 | 6580 | 7011 | 7011 | 7716 | 5q11.2 |
| MAP3K4 | ENSG00000085511 | 4502 | 5444 | 5490 | 7522 | 6q26 |
| MAPK1 | ENSG00000100030 | 4566 | 1487 | 11022 | 11336 | 22q11.22 |
| MAX | ENSG00000125952 | 1130 | 728 | 3155 | 4430 | 14q23.3 |
| MECOM | ENSG00000085276 | 3176 | 3593 | 5732 | 9103 | 3q26.2 |
| MED12 | ENSG00000184634 | 5978 | 6827 | 6977 | 7425 | Xq13.1 |
| MEN1 | ENSG00000133895 | 2399 | 2868 | 3162 | 4246 | 11q13 |
| MET | ENSG00000105976 | 4002 | 4632 | 6635 | 7039 | 7q31 |
| MGA | ENSG00000174197 | 9527 | 11859 | 14439 | 17651 | 15q15 |
| MGMT | ENSG00000170430 | 1113 | 867 | 1759 | 3557 | 10q26.3 |
| MLH1 | ENSG00000076242 | 1680 | 2230 | 2752 | 3532 | 3p22.2 |
| MSH2 | ENSG00000095002 | 2898 | 2918 | 3628 | 4552 | 2p21-p16.3 |
| MSH3 | ENSG00000113318 | 3229 | 4092 | 4092 | 4709 | 5q14.1 |
| MSH6 | ENSG00000116062 | 3506 | 4055 | 7476 | 10993 | 2p16.3 |
| MTOR | ENSG00000198793 | 5810 | 8677 | 8677 | 12119 | 1p36.22 |
| MUC6 | ENSG00000184956 | 6552 | 8006 | 8006 | 8428 | 11p15.5 |
| MYC | ENSG00000136997 | 1899 | 2150 | 2364 | 3001 | 8q24.21 |
| MYCN | ENSG00000134323 | 2602 | 2602 | 2602 | 2602 | 2p24.3 |
| MYD88 | ENSG00000172936 | 1795 | 1279 | 3314 | 3989 | 3p22.2 |
| MYH9 | ENSG00000100345 | 4234 | 7501 | 7501 | 8959 | 22q12.3 |
| NCOR1 | ENSG00000141027 | 5130 | 3490 | 10720 | 15404 | 17p12-p11.2 |
| NF1 | ENSG00000196712 | 7359 | 8223 | 12425 | 26685 | 17q11.2 |
| NF2 | ENSG00000186575 | 2924 | 2020 | 6025 | 7170 | 22q12.2 |
| NFE2L2 | ENSG00000116044 | 1569 | 979 | 2853 | 5907 | 2q31.2 |
| NIPBL | ENSG00000164190 | 7683 | 8729 | 10435 | 12529 | 5p13.2 |
| NOTCH1 | ENSG00000148400 | 8257 | 9371 | 9371 | 9880 | 9q34.3 |
| NOTCH2 | ENSG00000134250 | 6648 | 5041 | 11389 | 16804 | 1p12 |
| NPM1 | ENSG00000181163 | 1136 | 1237 | 1758 | 3828 | 5q35.1 |
| NRAS | ENSG00000213281 | 4449 | 4449 | 4449 | 4449 | 1p13.2 |
| NSD1 | ENSG00000165671 | 7014 | 7688 | 12892 | 15242 | 5q35.3 |
| NSD2 | ENSG00000109685 | 5300 | 7534 | 8568 | 19776 | 4p16.3 |
| NUP133 | ENSG00000069248 | 3804 | 5207 | 5207 | 6080 | 1q42.13 |
| NUP93 | ENSG00000102900 | 2475 | 2741 | 8257 | 11136 | 16q13 |
| PAX5 | ENSG00000196092 | 1724 | 1112 | 8615 | 9333 | 9p13.2 |
| PBRM1 | ENSG00000163939 | 4513 | 4905 | 7523 | 9654 | 3p21.1 |
| PCBP1 | ENSG00000169564 | 1750 | 1750 | 1750 | 1750 | 2p13.3 |
| PDGFRA | ENSG00000134853 | 3182 | 2417 | 6576 | 9732 | 4q12 |
| PDS5B | ENSG00000083642 | 5038 | 5246 | 7497 | 13212 | 13q13.1 |
| PGR | ENSG00000082175 | 3846 | 2633 | 13748 | 15306 | 11q22.1 |
| PHF6 | ENSG00000156531 | 3785 | 4489 | 4759 | 8015 | Xq26.2 |
| PIK3CA | ENSG00000121879 | 6359 | 9093 | 9093 | 9411 | 3q26.32 |
| PIK3CB | ENSG00000051382 | 3449 | 3181 | 5919 | 7850 | 3q22.3 |
| PIK3CG | ENSG00000105851 | 3930 | 5197 | 5377 | 5930 | 7q22.3 |
| PIK3R1 | ENSG00000145675 | 2734 | 2473 | 7011 | 10767 | 5q13.1 |
| PIK3R2 | ENSG00000105647 | 2410 | 2187 | 4033 | 4524 | 19p13.11 |
| PIM1 | ENSG00000137193 | 1698 | 1734 | 2650 | 2977 | 6p21.2 |
| PLCB4 | ENSG00000101333 | 5045 | 5558 | 5833 | 6900 | 20p12.3-p12.2 |
| PLCG1 | ENSG00000124181 | 2633 | 1005 | 5490 | 7784 | 20q12 |
| PLXNB2 | ENSG00000196576 | 3861 | 3443 | 6383 | 7257 | 22q13.33 |
| PMS1 | ENSG00000064933 | 2363 | 2795 | 3417 | 5448 | 2q32.2 |
| PMS2 | ENSG00000122512 | 2009 | 1719 | 2855 | 3936 | 7p22.1 |
| POLE | ENSG00000177084 | 5540 | 6800 | 8011 | 16609 | 12q24.33 |
| POLQ | ENSG00000051341 | 7760 | 8775 | 9055 | 9400 | 3q13.33 |
| POLRMT | ENSG00000099821 | 2013 | 1106 | 3836 | 5124 | 19p13.3 |
| PPM1D | ENSG00000170836 | 3194 | 2996 | 4778 | 5044 | 17q23.3 |
| PPP2R1A | ENSG00000105568 | 2207 | 1233 | 5380 | 7437 | 19q13.41 |
| PPP6C | ENSG00000119414 | 3409 | 4172 | 4349 | 4443 | 9q33.3 |
| PRKAR1A | ENSG00000108946 | 2038 | 1209 | 4327 | 6821 | 17q24.2 |
| PSIP1 | ENSG00000164985 | 2182 | 1966 | 3391 | 6768 | 9p22.3 |
| PTCH1 | ENSG00000185920 | 5580 | 7659 | 10631 | 16591 | 9q22.32 |
| PTEN | ENSG00000171862 | 3760 | 1605 | 9027 | 11662 | 10q23.31 |
| PTMA | ENSG00000187514 | 949 | 933 | 1635 | 3378 | 2q37.1 |
| PTPDC1 | ENSG00000158079 | 4202 | 4437 | 4517 | 6342 | 9q22.32 |
| PTPN11 | ENSG00000179295 | 3677 | 1876 | 6101 | 6996 | 12q24.13 |
| PTPRC | ENSG00000081237 | 2907 | 2336 | 5164 | 8371 | 1q31.3-q32.1 |
| PTPRD | ENSG00000153707 | 7128 | 8242 | 9911 | 11045 | 9p24.1-p23 |
| RAC1 | ENSG00000136238 | 1080 | 845 | 2323 | 3114 | 7p22.1 |
| RAD21 | ENSG00000164754 | 1698 | 896 | 3749 | 6192 | 8q24.11 |
| RAF1 | ENSG00000132155 | 2154 | 2354 | 3300 | 7847 | 3p25.2 |
| RARA | ENSG00000131759 | 1942 | 2024 | 3041 | 4731 | 17q21.2 |
| RASA1 | ENSG00000145715 | 3543 | 3752 | 4979 | 5232 | 5q14.3 |
| RB1 | ENSG00000139687 | 3198 | 4840 | 4840 | 6169 | 13q14.2 |
| RBM10 | ENSG00000182872 | 2972 | 3201 | 3747 | 3889 | Xp11.3 |
| RET | ENSG00000165731 | 4292 | 4337 | 5659 | 6661 | 10q11.21 |
| RFC1 | ENSG00000035928 | 2518 | 1255 | 4886 | 6713 | 4p14 |
| RHEB | ENSG00000106615 | 1064 | 900 | 2075 | 3535 | 7q36.1 |
| RHOA | ENSG00000067560 | 977 | 889 | 2031 | 2582 | 3p21.31 |
| RHOB | ENSG00000143878 | 2372 | 2372 | 2372 | 2372 | 2p24.1 |
| RIT1 | ENSG00000143622 | 1289 | 855 | 3445 | 3919 | 1q22 |
| RNF111 | ENSG00000157450 | 3832 | 4803 | 5913 | 7181 | 15q22.1-q22.2 |
| RNF43 | ENSG00000108375 | 3789 | 3963 | 5575 | 7505 | 17q23.2 |
| RPL22 | ENSG00000116251 | 960 | 693 | 2279 | 4900 | 1p36.31 |
| RPL5 | ENSG00000122406 | 906 | 1043 | 1210 | 2890 | 1p22.1 |
| RPS6KA3 | ENSG00000177189 | 3595 | 847 | 7918 | 8837 | Xp22.12 |
| RRAS2 | ENSG00000133818 | 1081 | 970 | 2034 | 3453 | 11p15.2 |
| RUNX1 | ENSG00000159216 | 2760 | 1590 | 7274 | 15574 | 21q22.12 |
| RXRA | ENSG00000186350 | 4133 | 5215 | 5770 | 10814 | 9q34.2 |
| SCAF4 | ENSG00000156304 | 3524 | 3846 | 4193 | 5901 | 21q22.11 |
| SETBP1 | ENSG00000152217 | 5671 | 5842 | 9899 | 11151 | 18q12.3 |
| SETD2 | ENSG00000181555 | 6928 | 7555 | 8172 | 10019 | 3p21.31 |
| SF1 | ENSG00000168066 | 2176 | 2778 | 3470 | 5972 | 11q13.1 |
| SF3B1 | ENSG00000115524 | 3112 | 2003 | 6526 | 9164 | 2q33.1 |
| SIN3A | ENSG00000169375 | 3706 | 4930 | 6737 | 12453 | 15q24.2 |
| SMAD2 | ENSG00000175387 | 8209 | 1842 | 34526 | 36366 | 18q21.1 |
| SMAD4 | ENSG00000141646 | 3392 | 1371 | 8769 | 13123 | 18q21.2 |
| SMARCA1 | ENSG00000102038 | 3770 | 3948 | 4291 | 4363 | Xq25-q26.1 |
| SMARCA4 | ENSG00000127616 | 4722 | 5193 | 5691 | 11067 | 19p13.2 |
| SMARCB1 | ENSG00000099956 | 1408 | 1643 | 1728 | 2481 | 22q11.23 |
| SMC1A | ENSG00000072501 | 7608 | 9784 | 9930 | 10484 | Xp11.22 |
| SMC3 | ENSG00000108055 | 3323 | 4114 | 4114 | 4275 | 10q25.2 |
| SOS1 | ENSG00000115904 | 5884 | 8314 | 8517 | 9263 | 2p22.1 |
| SOX17 | ENSG00000164736 | 2342 | 2342 | 2342 | 2342 | 8q11.23 |
| SOX9 | ENSG00000125398 | 3935 | 3935 | 3935 | 3935 | 17q24.3 |
| SPOP | ENSG00000121067 | 1307 | 919 | 2985 | 6380 | 17q21.33 |
| SPTA1 | ENSG00000163554 | 6500 | 7998 | 7999 | 10025 | 1q23.1 |
| SPTAN1 | ENSG00000197694 | 7118 | 7857 | 7872 | 11006 | 9q34.11 |
| SRSF2 | ENSG00000161547 | 1279 | 1357 | 2885 | 3017 | 17q25.2 |
| STAG2 | ENSG00000101972 | 3733 | 4393 | 6045 | 8187 | Xq25 |
| STK11 | ENSG00000118046 | 2075 | 2611 | 3554 | 8639 | 19p13.3 |
| TAF1 | ENSG00000147133 | 4265 | 2295 | 7722 | 10860 | Xq13.1 |
| TBL1XR1 | ENSG00000177565 | 2517 | 604 | 7948 | 10614 | 3q26.32 |
| TBX3 | ENSG00000135111 | 3297 | 4208 | 4723 | 5950 | 12q24.21 |
| TCF12 | ENSG00000140262 | 2592 | 1809 | 6061 | 9446 | 15q21.3 |
| TCF7L2 | ENSG00000148737 | 2973 | 3953 | 4136 | 4935 | 10q25.2-q25.3 |
| TENT5D | ENSG00000174016 | 3070 | 3106 | 3106 | 3204 | Xq21.1 |
| TET2 | ENSG00000168769 | 8083 | 9679 | 10166 | 16474 | 4q24 |
| TGFBR2 | ENSG00000163513 | 4612 | 4605 | 4621 | 4704 | 3p24.1 |
| TGIF1 | ENSG00000177426 | 1220 | 1088 | 3030 | 8222 | 18p11.31 |
| THRAP3 | ENSG00000054118 | 2964 | 3339 | 4432 | 4736 | 1p34.3 |
| TLR4 | ENSG00000136869 | 3338 | 3908 | 4844 | 5256 | 9q33.1 |
| TMSB4X | ENSG00000205542 | 824 | 628 | 1702 | 1705 | Xp22.2 |
| TNFAIP3 | ENSG00000118503 | 3022 | 3817 | 4735 | 5051 | 6q23.3 |
| TP53 | ENSG00000141510 | 2045 | 2506 | 2724 | 3936 | 17p13.1 |
| TRAF3 | ENSG00000131323 | 3391 | 2239 | 7700 | 8263 | 14q32.32 |
| TSC1 | ENSG00000165699 | 5935 | 8310 | 8604 | 9911 | 9q34 |
| TSC2 | ENSG00000103197 | 4421 | 5287 | 6156 | 11121 | 16p13.3 |
| TXNIP | ENSG00000265972 | 1876 | 1509 | 2926 | 3625 | 1q21.1 |
| U2AF1 | ENSG00000160201 | 1561 | 966 | 4694 | 11112 | 21q22.3 |
| UNCX | ENSG00000164853 | 2048 | 2048 | 2048 | 2048 | 7p22.3 |
| USP9X | ENSG00000124486 | 8988 | 8371 | 12401 | 13962 | Xp11.4 |
| VHL | ENSG00000134086 | 2434 | 2696 | 3737 | 4213 | 3p25.3 |
| WT1 | ENSG00000184937 | 2210 | 2466 | 3122 | 4113 | 11p13 |
| XPO1 | ENSG00000082898 | 3166 | 3479 | 6776 | 11708 | 2p15 |
| ZBTB20 | ENSG00000181722 | 1709 | 635 | 3740 | 8279 | 3q13.31 |
| ZBTB7B | ENSG00000160685 | 2398 | 3594 | 4017 | 4303 | 1q21.3 |
| ZC3H12A | ENSG00000163874 | 1808 | 2684 | 2684 | 3117 | 1p34.3 |
| ZCCHC12 | ENSG00000174460 | 2231 | 2231 | 2231 | 2231 | Xq24 |
| ZFHX3 | ENSG00000140836 | 12460 | 13841 | 16064 | 17503 | 16q22.2-q22.3 |
| ZFP36L1 | ENSG00000185650 | 1868 | 1876 | 3190 | 5160 | 14q24.1 |
| ZFP36L2 | ENSG00000152518 | 2316 | 1627 | 3696 | 3696 | 2p21 |
| ZMYM2 | ENSG00000121741 | 6572 | 5445 | 10247 | 13618 | 13q12.11 |
| ZMYM3 | ENSG00000147130 | 5104 | 5723 | 6067 | 6916 | Xq13.1 |
| ZNF133 | ENSG00000125846 | 2093 | 2622 | 4060 | 5399 | 20p11.23 |
| ZNF750 | ENSG00000141579 | 2613 | 2613 | 3713 | 3713 | 17q25.3 |

**References**

Bailey MH, Tokheim C, Porta-Pardo E, Sengupta S, Bertrand D, Weerasinghe A, Colaprico A, Wendl MC, Kim J, Reardon B, et al. 2018. Comprehensive Characterization of Cancer Driver Genes and Mutations. Cell 173:371-385 e318.

Grimmett G, Stirzaker D. 2009. Probability and random processes. Oxford; New York: Oxford University Press.

Lawrence MS, Stojanov P, Mermel CH, Robinson JT, Garraway LA, Golub TR, Meyerson M, Gabriel SB, Lander ES, Getz G. 2014. Discovery and saturation analysis of cancer genes across 21 tumour types. Nature 505:495-501.

Wood RD, Mitchell M, Lindahl T. 2005. Human DNA repair genes, 2005. Mutat Res 577:275-283.

Wood RD, Mitchell M, Sgouros J, Lindahl T. 2001. Human DNA repair genes. Science 291:1284-1289.
